# Proteogenomic analysis reveals RNA as an important source for tumor-agnostic neoantigen identification correlating with T-cell infiltration

**DOI:** 10.1101/2022.09.17.508207

**Authors:** Celina Tretter, Niklas de Andrade Krätzig, Matteo Pecoraro, Sebastian Lange, Philipp Seifert, Clara von Frankenberg, Johannes Untch, Florian S Dreyer, Eva Bräunlein, Mathias Wilhelm, Daniel P Zolg, Thomas Engleitner, Sebastian Uhrig, Melanie Boxberg, Katja Steiger, Julia Slotta-Huspenina, Sebastian Ochsenreither, Nikolas von Bubnoff, Sebastian Bauer, Melanie Boerries, Philipp J Jost, Kristina Schenck, Iska Dresing, Florian Bassermann, Helmut Friess, Daniel Reim, Konrad Grützmann, Katrin Pfütze, Barbara Klink, Evelin Schrock, Bernhard Haller, Bernhard Kuster, Matthias Mann, Wilko Weichert, Stefan Fröhling, Roland Rad, Michael Hiltensperger, Angela M Krackhardt

**Author notes:** Corresponding author: Angela M. Krackhardt, Klinik und Poliklinik für Innere Medizin III, Technische Universität München, School of Medicine, Ismaningerstr. 22, 81675 München. These authors contributed equally to this work (Celina Tretter, Niklas de Andrade Krätzig). These authors jointly supervised this work (Michael Hiltensperger, Angela M Krackhardt).

## Abstract

Systemic pan-tumor analyses may reveal the significance of common features implicated in cancer immunogenicity and patient survival. Here, we provide a comprehensive multi-omics data set for 32 patients across 25 tumor types by combining proteogenomics with phenotypic and functional analyses. By using an optimized computational approach, we discovered a large number of novel tumor-specific and tumor-associated antigens including shared common target candidates. To create a pipeline for the identification of neoantigens in our cohort, we combined deep DNA and RNA sequencing with MS-based immunopeptidomics of tumor specimens, followed by the assessment of their immunogenicity. In fact, we could detect a broad variety of non-wild type HLA-binding peptides in the majority of patients and confirmed the immunogenicity of 24 neoantigens. Most interestingly, the majority of total and immunogenic neoantigens originated from variants identified in the RNA dataset, illustrating the importance of RNA as a still understudied source of cancer antigens. Moreover, the amount of these mainly RNA-based immunogenic neoantigens correlated positively with overall CD8^+^ tumor-infiltrating T cells. This study therefore underlines the importance of RNA-centered variant detection for the identification of shared biomarkers and potentially relevant neoantigen candidates.

**Statement of significance:** The significance of this study lies not only in the potential of our optimized proteogenomic workflow for the discovery of neoantigens (in particular RNA-derived neoantigens) for clinical application, but sheds light on the entity-agnostic prevalence of HLA class I peptide presentation of RNA processing events to be used for tumor targeting.

## Introduction

Genetic aberrations are not only centrally involved in the development of cancer but may also result in the formation of neoantigens that have the potential to mount an anti-tumor immune response. Such neoantigens can be recognized as foreign and targeted by neoantigen-specific T cells. Thus, the identification of such neoantigens is becoming increasingly important for the development of novel immunotherapies (1–5). However, the vast majority of neoantigens are not shared between cancer patients and the validation of in silico-predicted neoantigen candidates that range in the thousands is often limited or impractical in a clinical setting. For this reason, our group reported a proteogenomic approach that combines mass spectrometry (MS) of immunoprecipitated HLA class I (pHLA-I) peptides with whole exome sequencing (WES) of melanoma tumors for the identification and validation of such neoantigens at the protein level (6). We were able to show for the first time that such a proteogenomic approach is feasible in fresh solid tumor material and yields a refined number of immunogenic neoantigens. Yet the number of neoantigens that could be identified with our approach was limited and the findings had to be validated in different cancer entities.

It was reported that not only somatic mutations on coding exons represent a source of neoantigens but also non-coding transcripts, intronic regions and splice sites (7–10). Furthermore, RNA processing events such as RNA editing have been investigated in more detail lately. RNA editing is a widespread post-transcriptional mechanism conferring specific and reproducible nucleotide changes in selected RNA transcripts that occurs in normal cells (11) but is also involved in disease pathogenesis and is altered in cancer (12–14). These events have been recently associated with diversifying the cancer proteome (14, 15) and RNA variants derived from editing events were further investigated in more detail as a source of aberrantly expressed peptides (16, 17). As RNA regulation is mediated by *cis* regulatory elements and *trans* regulatory factors which are often disrupted by somatic mutations or affected by oncogenic signaling (18), antigens derived from cancer-associated RNA editing may represent in part true neoantigens and are therefore of high interest for targeted cancer immunotherapy. Thus, we included tumor transcriptomics in addition to WES, to detect neoantigens that were derived from RNA processing events.

Furthermore, we previously showed that integrating spectral prediction features into the MS-spectra matching process during neoantigen identification, known as rescoring, is a powerful method to deal with larger search spaces and it increases coverage and sensitivity of the analysis (19, 20). Therefore, we added the artificial intelligence algorithm Prosit and utilized a Prosit-based rescoring workflow in our pipeline for neoantigen identification (20, 21).

In this study, we use a subset of 32 patients with different tumor entities that were mainly included in the previously described MASTER cohort (22) to test our improved proteogenomic pipeline in a cross-entity cohort ImmuNEO MASTER. We discover many shared genetic variants and tumor-associated peptides between patients independent of the tumor entity. Most importantly, in the majority of patients we identify neoantigens that were predominantly derived from RNA sources. In addition, we perform T cell phenotyping in the tumor microenvironment and show that immunogenic neoantigens correlate with increased CD8^+^ T-cell infiltration. Thus, these data demonstrate that proteogenomic-based neoantigen identification is feasible in a cross-entity cohort and that neoantigens originating from RNA sources might present highly relevant targets for the development of novel immunotherapies.

## Results

This study took advantage of a patient cohort included in the MASTER Program (22). Detailed information about patient samples and respective analyses are described in the Methods section and are listed in Suppl. Table S1A and B.

For the identification of common tissue-agnostic immune-related hallmarks and neoantigen candidates in our cross-entity cohort ImmuNEO MASTER (Suppl. Table S1A, B and Suppl. Figure S1A, B), we created a general workflow for the analyses of tumor specimens which is illustrated in Figure 1. First, tumor-infiltrating immune cells were characterized in the tumor microenvironment (TME) of fresh tumor tissue by flow cytometric immunophenotyping as well as transcriptome analyses of sorted CD8^+^ T cells. Next, for the respective characterization of indicated tumor specimens we used WES/whole genome sequencing (WGS) and RNA sequencing (RNA-seq) data from patients included in the MASTER cohort or from the ImmuNEO Plus samples that were respectively analyzed at the same DKFZ facility as the samples of the MASTER cohort (22). The analytical core of our neoantigen discovery pipeline is its proteogenomic approach. For this, we performed immunoprecipitation of pHLA-I with subsequent MS analysis for the identification of the presented immunopeptidome. We then used an optimized workflow of our previously published strategy (6) for the identification of neoantigens by combining the personalized genomic data with the MS-based immunopeptidomic data using pFIND (23). As critical innovations we included RNA-seq data and used the artificial intelligence algorithm Prosit for increased coverage and sensitivity of our neoantigen discovery pipeline (20, 21). Immunogenicity of the identified neoantigen candidates was assessed *in vitro* by using patient-derived autologous or healthy donor (HD)-derived allogenic-matched T cells. Finally, in order to decipher potential clinical conditions for the identification of neoantigens which might be crucial knowledge for clinical application, we correlated the number of identified total and immunogenic neoantigens with the TME immunophenotyping data.

**Figure 1.**
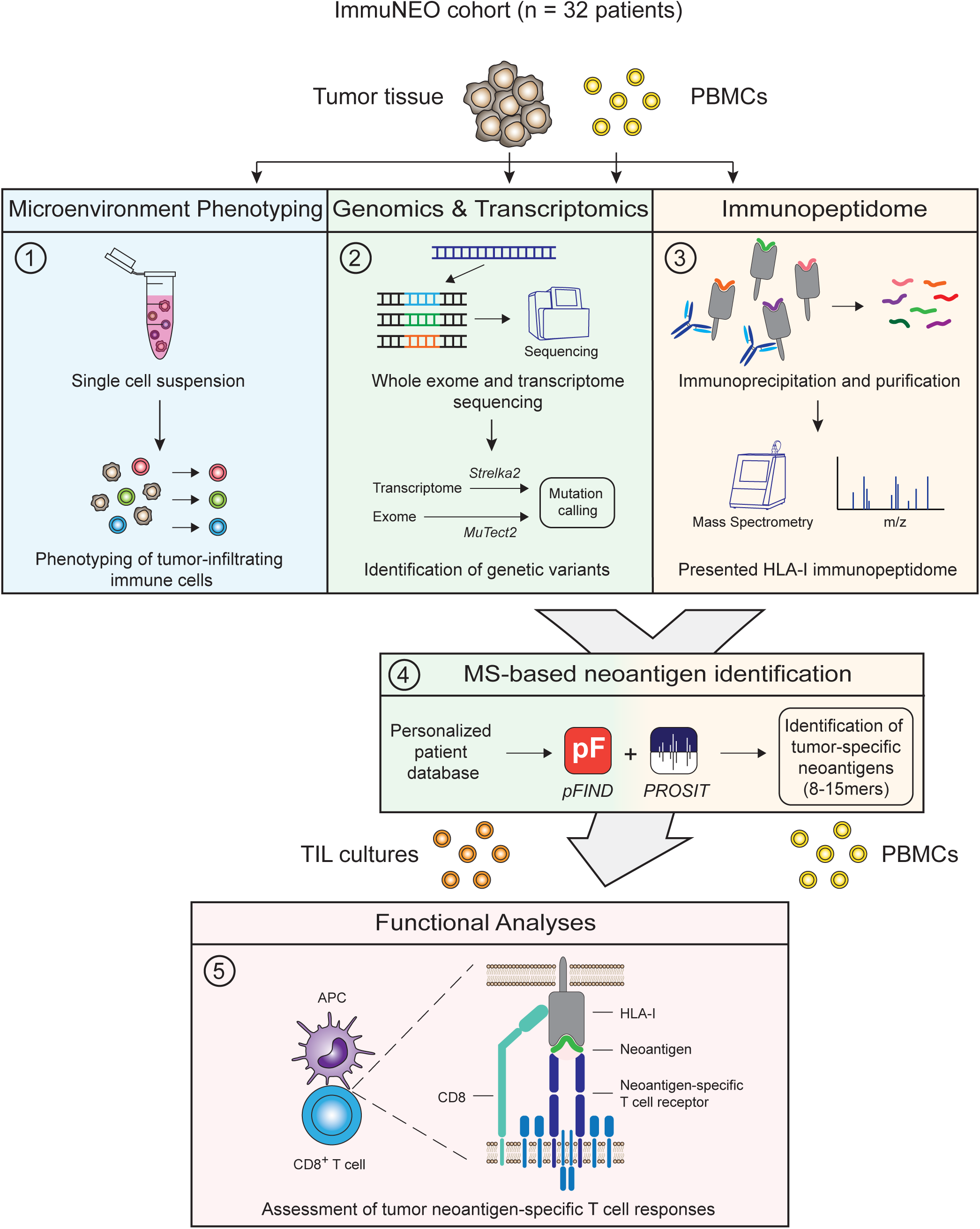
Overview of the workflow for immunophenotyping, proteogenomic and functional analyses for neoantigen identification in the cross-entity cohort. Tumor material and peripheral blood from 32 patients included into the ImmoNEO MASTER cohort harboring diverse tumor entities was used for the following analyses: (1) tumor microenvironment phenotyping; fresh primary tumor tissue was enzymatically digested and single cells were used for multi-colour flow cytometric analysis of several immune cells and phenotypic markers. In parallel, FACS-sorted CD8^+^ T cells were used for bulk transcriptome analysis (RNA-seq). (2) Genomic and transcriptomic analysis; primary tumor tissue was used for whole exome (WES)/whole genome sequencing (WGS) and RNA-seq. Blood from the same patient served as control samples for WES/WGS analyses. Mutations were called by MuTect2 (v4.1.0.0) from WES/WGS data and by Strelka2 (v2.9.10) from RNA-seq data and mutations were filtered for short nucleotide polymorphisms (SNPs) by using the dbSNP database. 3) Immunopeptidome analysis; fresh primary tumor tissue was used for HLA class I-bound peptide immunoprecipitation and subsequent mass spectrometry (MS) analysis of eluted peptides. The whole HLA class I peptidome was analysed using pFIND (v3.1.5) with 1% FDR on the spectral level looking for 8-15mers. (4) MS-based neoantigen identification; patient-specific mutational data from (2) were used to generate a personalized database. Therefore, all full length mutated ORFs generated by VCF-translate (v1.5) were added to the Ensembl92 data set and matched with the MS-identified peptide sequences using pFIND with 5% FDR on the spectral level looking for 8- 15mers. By filtering for peptides only matching to the mutated ORF sequences, tumor-specific neoantigen candidates were identified. The machine learning tool Prosit was additionally integrated to rescore the peptide spectra matching to the patient-specific ORF database. Afterwards several filtering and post-processing steps were applied for the identification of neoantigen candidates. (5) Immunogenicity assessment of neoantigen candidates; patient-derived autologous immune cells (PBMCs and TILs) as well as selected allogenic-matched healthy donor-derived PBMCs were tested for immunogenicity in response to the identified neoantigen candidates using a modified accelerated co-cultured dendritic cell (acDC) protocol to identify immunogenic neoantigens. APC, antigen-presenting cell; FDR, false discovery rate; HLA-I, human leukocyte antigen class I; ORF, open reading frame; PBMC, peripheral blood mononuclear cells; TIL, tumor-infiltrating lymphocytes.

### The phenotype of tumor-infiltrating T cells is independent of the tumor entity

To study if we could observe tumor-agnostic immunological features in the immune TME and correlate them with clinical outcome, we performed flow cytometric immunophenotyping of fresh primary tumor tissues. In 17 patients, from whom enough tumor material was available, T cell subsets were examined.

First, we looked at the relative cell numbers of CD8^+^ T cells per gram tumor (Figure 2A). The two melanoma specimens and the pancreatic cancer metastasis of a patient with mismatch repair deficiency (dMMR) (ImmuNEO-11 T2) demonstrated a high amount of T-cell infiltration matching to the high mutational burden often present in these malignancies (24, 25). However, also other tumor entities, including a sarcoma specimen (ImmuNEO-5), showed high amounts of tumor-infiltrating lymphocytes (TILs) (Figure 2A). CD8^+^ and CD4^+^ T cells predominantly consisted of effector memory T (Tem; CD45RA^-^CD62L^low^) cells regardless of the tumor entity (Figure 2B and Suppl. Figure S2A, B). Moreover, the distribution of CD8^+^ T cell subsets and – to a lesser extent – of CD4^+^ T cell subsets between different metastases of a defined individual patient were highly comparable independent of their anatomical metastatic location (Figure 2B and Suppl. Figure S2B) and despite differences in their relative cell numbers (Figure 2A). Since the functional state of TILs is linked to their potential anti-tumor activity, we analyzed the expression of selected activation markers (HLA-DR and CD103) and inhibitory markers (PD-1, TIM-3, and LAG-3). To account for differences in overall cell numbers and to investigate the activation status on a population level, we looked into the frequencies of activation or inhibitory markers on CD8^+^ and CD4^+^ T cells (Suppl. Figure S2C), respectively, that express at least one marker. There was no difference in the frequencies of CD8^+^ T cells with activation markers between different tumor entities, and tumor specimens with high frequencies of inhibitory markers were present in carcinoma, sarcoma, and melanoma patients (Figure 2C).

**Figure 2.**
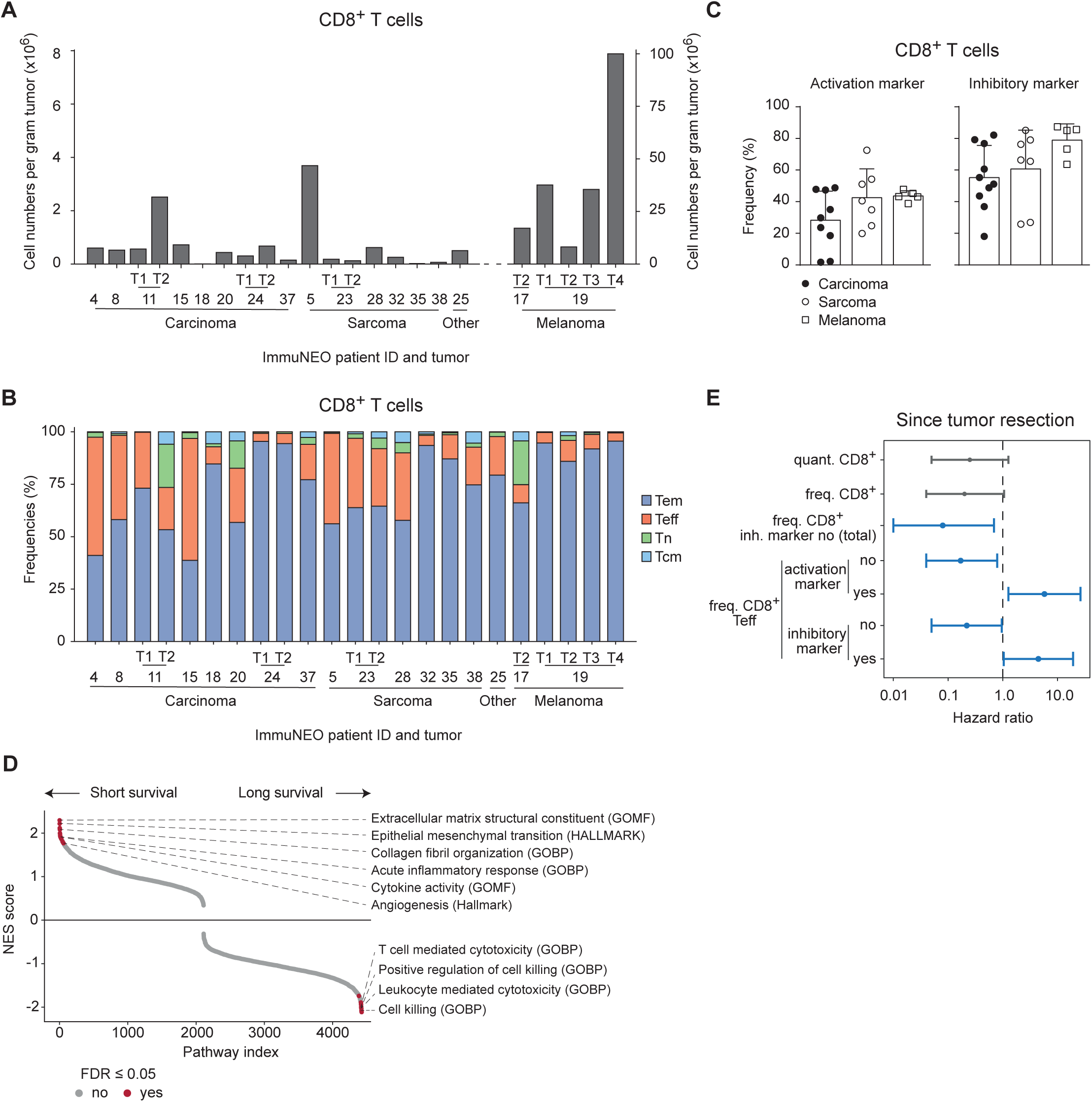
Phenotypic and transcriptomic investigation of the immune tumor microenvironment of a defined subgroup of the ImmuNEO MASTER cohort. **A,** Quantitative numbers of CD8^+^ T cells per gram tumor identified by flow cytometric assessment of fresh tumor tissue per patient grouped by tumor entity. **B,** Frequencies of different CD8^+^ T cell subsets of all identified tumor infiltrating CD8^+^ T cells per patient grouped by tumor entity. **C,** Frequencies of CD8^+^ T cells expressing at least one activation marker (HLA-DR, CD103) or inhibitory marker (PD-1, TIM- 3, LAG-3) for different cancer entities. Symbols depict individual tumor samples. Data are shown as mean + s.d.. **D**, Gene set enrichment analysis (GSEA) for gene signatures differentially expressed in sorted tumor-infiltrating CD8^+^ T cells from bulk RNA sequencing (RNA-seq) of patients with short (below 1 year, n = 3) and long survival (above 1 year, n = 5) since tumor resection. NES scores for each pathway are depicted and significantly enriched (p ≤ 0.05) pathways are coloured in red. **E,** Forest plot showing the hazard ratio (dot) and 95% confidence intervals (lines) calculated by log rank test and Cox’s proportional hazards model of several phenotypic parameters for the survival of patients since tumor resection (n = 17). Significant correlations (p ≤ 0.05) are highlighted in blue. For statistical analysis only one representative tumor sample per patient was used (see core cohort Suppl. Table S1A). **A, B,** n = 23 tumor samples from n = 17 patients (see Suppl. Table S1A). **C,** n = 22 tumor samples from n = 16 patients. FDR, false discovery rate; freq., frequency; GOBP, Gene ontology biological function gene set; GOMF, Gene ontology molecular function gene set; HALLMARK, hallmark gene set; inh., inhibitory; NES, normalized enrichment score; quant., quantified per gram tumor; T, tumor; Tcm, central memory T cells; Teff, effector T cells; Tem, effector memory T cells; Tn, naïve T cells.

In order to identify clinically relevant transcriptional T cell signatures in our cohort, we performed RNA-seq on sorted CD8^+^ T cells from eight patients. Patients were grouped based on their survival data since tumor resection into a short survival (less than 1 year) and a long survival (more than 1 year) group (Suppl. Figure S2D and Suppl. Table S1A). By using gene set enrichment analyses (GSEA), we could show that pathways associated with T cell-mediated cytotoxic functions were upregulated in the long survival group, while pathways associated with general inflammatory responses were upregulated in the short survival group (Figure 2D). In addition, to identify tissue-agnostic features that correlate with survival, the influence of each parameter on the survival of our patients since tumor resection was assessed by log rank test and Cox’s proportional hazards model (Figure 2E, Suppl. Figure S2E). Although the quantified numbers and frequencies of CD8^+^ T cells showed only a non-significant trend for a positive correlation with increased survival, the overall frequency of CD8^+^ T cells without inhibitory markers in the TME correlated positively with increased survival (Figure 2E). Moreover, the frequencies of cells without activation or inhibitory markers within the CD8^+^ Teff subset correlated positively as well with increased survival and, consequently, a high fraction of cells with activation or inhibitory markers within this subset correlated positively with reduced survival (Figure 2F). Of note, we observed only non-significant trends for CD4^+^ T cells (Suppl. Figure S2E).

In summary, we observed that tumor-infiltrating T cells in our heterogenous pan-cancer cohort were mainly comprised of Tem cells independent of the tumor entity. Moreover, we could reproduce findings that had previously been observed in homogenous tumor cohorts, such as increased numbers of TILs in malignancies that are characterized by high mutational burden, and observed specific transcriptional pathways in CD8^+^ T cells that were associated with clinical outcome (26) in this cross-entity cohort.

### Genetic variants are more common at the RNA level and are often shared between different tumor entities

In a next step, we assessed the number of genetic variants in the tumors at the DNA and RNA level. Since these data are the basis for the identification of neoantigen candidates and will later be cross-validated by our MS-based analyses of the tumor immunopeptidomes (Figure 1), we decided to use the datasets with unfiltered genetic variants to avoid loss of potential candidates (Suppl. Figure S3). Of note, the majority of genetic variants passed the filtering criteria at the RNA level for all tumor specimens but there were multiple exceptions regarding mutations at the DNA level.

The number of DNA and RNA variants varied greatly between patients but showed no clear deviation between different tumor entities in our pan-cancer cohort (Figure 3A). On average, we identified 302 somatic mutations per tumor, but a much higher number of genetic variants were identified at the RNA level, with an average of 4024 genetic variants per tumor (Figure 3A). Of note, the majority of DNA variants were also found at the RNA level (Suppl. Figure S4A), highlighting the power of RNA as a source for the discovery of genetic variants. In general, single-nucleotide substitutions accounted for most of the variants found at the DNA and RNA level but deletions and insertions as well as multi-nucleotide substitutions were also observed for some variants (Suppl. Figure S4B). Interestingly, there was no correlation between the number of DNA and RNA variants that were identified for each tumor (Suppl. Figure S4C), indicating that tumors with low levels of somatic mutations can still harbor a high amount of RNA variants.

**Figure 3.**
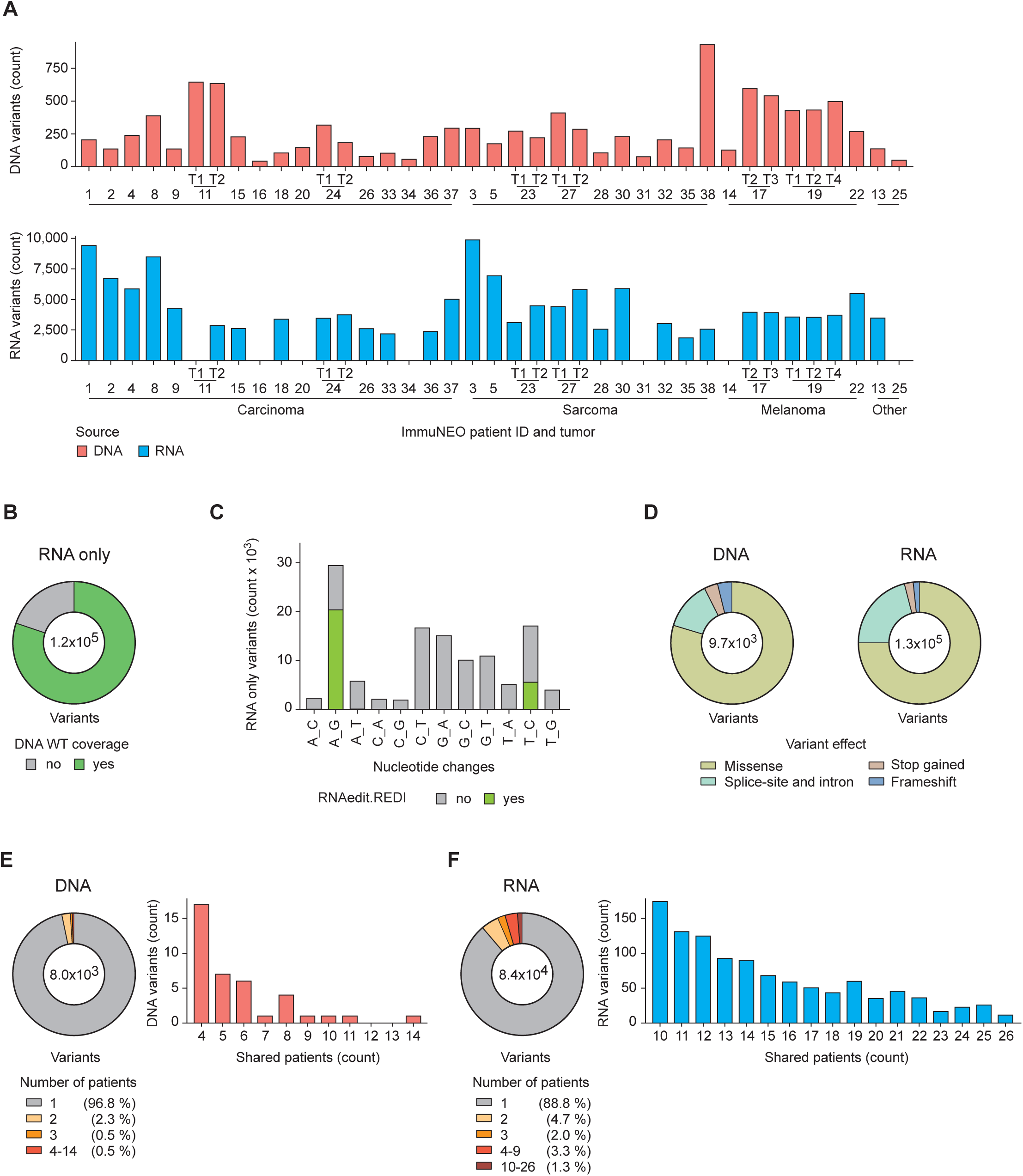
Genetic variants identified at the DNA and RNA level in tumor tissue from different cancer entities. **A,** Distribution of the total numbers of variants identified from DNA (upper panel) and RNA data (lower panel) identified per tumor sample grouped by tumor entity. Mutations were called by MuTect2 (v4.1.0.0) from whole exome (WES)/whole genome sequencing (WGS) data and by Strelka2 (v2.9.10) from RNA sequencing (RNA-seq) data. SNP-filtering was performed using the dbSNP-all data base. No RNA data was available for patients IN-11-T1, IN-14, IN-16, IN-20, IN-25, IN-31, IN-34. **B,** Pie chart depicting the proportion of variants only identified from RNA-seq data of all tumor samples combined where the respective wild type (WT) sequence was identified at the DNA level with a coverage of ≥ 3 reads (green) or the respective region was not covered at the DNA level (grey, < 3 reads). **C,** Distribution of the nucleotide exchange pattern over all single nucleotide variants only identified from RNA-seq data of all tumor samples combined. Variants previously identified in the REDIportal (29) database as RNA editing events are highlighted in green. **D,** Pie charts depicting the distribution of each mutation type for variants called from all DNA (left) and RNA (right) variants. **E, F,** Pie charts showing the proportions of unique and shared DNA variants (**E**) and RNA variants (**F**) between different patients. The right bar graph shows the number of variants shared by 4 to 14 patients for DNA variants (**E**) and shared by 10 to 26 patients for RNA variants (**F**) in more detail. **A-E,** n = 39 tumor samples from n = 32 patients for WES/WGS data; n = 32 tumor samples from n = 26 patients for RNA-seq data (see Suppl. Table S1A). T, tumor; WT, wild type.

The higher number of variants that were detected at the RNA level compared to the DNA level could be explained in part by more non-coding sources for RNA variants, such as regulatory RNAs and pseudogenes (Suppl. Figure S4D). However, these additional non-coding sources still did not account for this striking difference since most RNA variants were detected from protein coding regions (Suppl. Figure S4D). RNA editing events could present an additional source for RNA variants (11, 27). For this, we analyzed the coverage of the corresponding wild type (WT) locus at the DNA level and nucleotide exchange patterns for all variants that were only identified at the RNA level. Indeed, for most RNA variants we could detect a corresponding WT sequence at the DNA level (Figure 3B), suggesting that part of these variants might be derived from RNA editing events. In fact, a considerable portion of RNA variants harbored an adenosine (A) to guanosine (G) nucleotide exchange, which has been described in the context of RNA editing events (defined by A to inosine (I) editing, where I appears as G in RNA-seq data (28)) (11, 14), and the majority of variants with this specific nucleotide exchange have been reported as RNA editing events in the databank REDIportal (29) (Figure 3C). We observed that both DNA and RNA variants were mainly comprised of missense variants, but RNA variants consisted of more splice-site and intron variants (Figure 3D). Although the correlation between tumor mutational burden (TMB) (DNA variants per Mb) and increased survival was not statistically significant, we observed a positive trend and the overall number of DNA variants correlated positively with increased survival in our heterogenous cohort (Suppl. Figure S4E). There was no correlation between the number of genetic variants that were found solely at the RNA level and overall survival (Suppl. Figure S4E), suggesting that the sheer quantity of RNA variants does not present a prognostic biomarker for immunogenicity-associated survival.

Moreover, shared genetic mutations within this pan-cancer cohort were of special interest to us as these might lead to potential common neoantigens that could be attractive targets for immunotherapy. Therefore, we investigated in how many patients each genetic variant was detected. As expected, the vast majority of genetic variants were found to be unique at the DNA and RNA level (Figure 3E, F). Indeed, approximately 97% of variants were unique in our cohort at the DNA level (Figure 3E) but only 89% at the RNA level (Figure 3F). Together with the fact that we detected roughly 10 times more RNA variants compared to DNA variants, this means that we could identify approximately 37 times more shared genetic variants (detected in at least 2 patients) at the RNA level. In addition, we observed that a subset of RNA variants was shared in all patients, however, DNA variants were shared significantly less frequently and in smaller groups of patients (Figure 3E, F and Suppl. Table S2A, B).

To elucidate if these shared RNA variants were overlapping with each other in the same sets of patients, we focused on RNA variants that were found in at least ten tumor specimens with a minimum of two shared RNA variants (Suppl. Figure S4F). Overlapping shared RNA variants were not only commonly present in tumor metastases but also in different tumor entities in our pan-cancer cohort (Suppl. Figure S4F). Although the majority of shared RNA variants in these sets were found to be exclusive, we were able to identify 59 shared variants that showed some degree of overlap. Out of these, 11 RNA variants were present in all patients and tumor metastases of our pan-cancer cohort (Suppl. Tables S2B, C).

Taken together, we identified remarkably more genetic variants at the RNA level in general and shared variants in particular, and a substantial part of additional RNA variants was likely derived from RNA editing events.

### The tumor immunopeptidomes harbor many shared cancer-associated peptides across different tumor entities

To characterize the tumor immunopeptidomes in our pan-cancer cohort, we performed immunoprecipation of pHLA-I followed by MS analysis as previously described (6). Similar to the numbers of genetic variants, the overall numbers of peptides varied greatly between patients without a clear deviation between different tumor entities (Figure 4A and Suppl. Figure S5A). On average, approximately 5075 peptides could be identified per tumor (Figure 4A), with a length of 8 to 15 amino acids that were predominated by nonamers (Suppl. Figure S6). Exemplified in four patients (ImmuNEO- 4, -11, -14, -38), we analyzed the HLA anchor residues of the immunopeptides in all patients and could show that they were characteristic for the patients’ HLA composition with a purity of at least 95% (Suppl. Figure S7).

**Figure 4.**
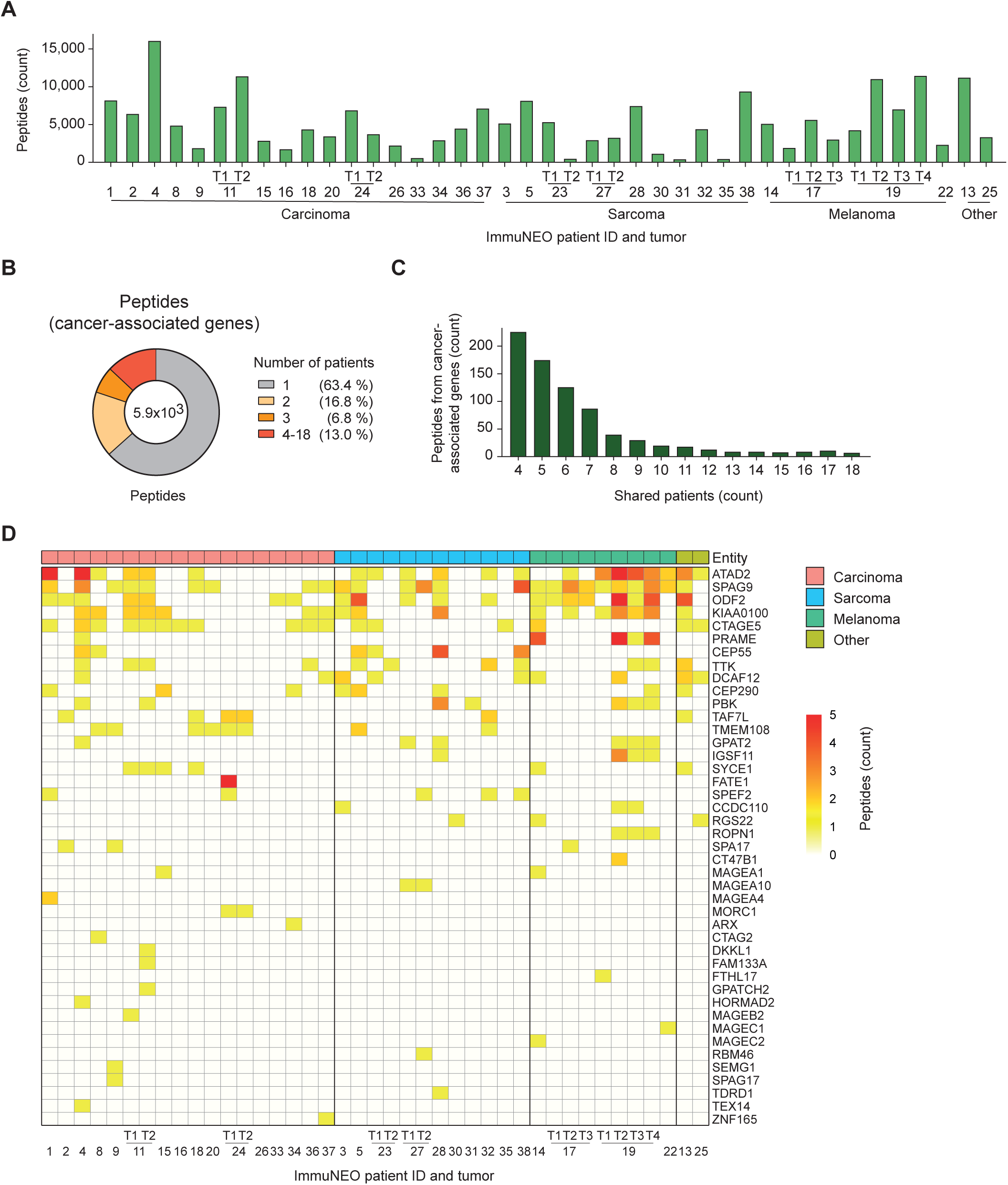
Analysis of the HLA class I tumor immunopeptidomes. **A,** Distribution of the total number of unique HLA class I peptides identified per tumor sample grouped by tumor entity. Peptides bound to HLA class I molecules on the surface of tumor cells were isolated by immunoprecipitation and sequenced by liquid chromatography with tandem mass spectrometry (LC-MS/MS). Peptide sequences were then mapped with 1% FDR to the Ensemble92 protein database using pFIND (v3.1.5) and unique sequences have been filtered. **B,** Pie chart showing the proportion of unique and shared peptides originating from cancer-associated genes (ProteinAtlas) between patients. **C,** Bar graph depicting the number of peptides shared by 4 to 18 patients in more detail. **D,** Heatmap depicting the numbers of unique peptides found per cancer testis antigen (CTA) gene in each tumor sample. Genes were sorted by the total number of peptides identified over all patients and samples were grouped by entity. **A-D,** n = 41 tumor samples from n = 32 patients (see Suppl. Table S1A). FDR, false discovery rate; HLA, human leukocyte antigen; T, tumor.

By focusing on peptides derived from cancer-associated genes that have been described in the Human Protein Atlas (30), we spotted that 36% of these peptides were shared between patients (Figure 4B) and a considerable number of them were present in up to 18 patients (Figure 4C). Out of these, 79 shared peptides showed some degree of overlap in at least eight tumor specimens (Suppl. Figure S5B). Moreover, 18 shared peptides were identified in at least 11 patients (Suppl. Figure S5B arrows and Suppl. Table S3) and were predicted by NetMHC4.0 to bind with good affinities to the patients’ HLA molecules (HLA-A03:01 or HLA-A11:01; Suppl. Table S1C). These peptide ligands have been previously described by several studies in the context of cancer (studies found on *PeptideAtlas*, 2022; *IEDB.org: Free epitope database and prediction resource*, 2022).

In addition, we analyzed peptides derived from reported cancer testis antigens (CTAs) using the CTpedia database (33) and discovered numerous CTA peptides in our cohort (Figure 4D). Although the majority of CTA peptides were only found to be unique in one patient, we identified multiple peptides derived from CTA-associated genes that were present in a substantial portion of patients independent of the tumor entity (e.g. ATAD2, SPAG9, ODF2, KIAA0100) (Figure 4D). Importantly, there was not only an overlap between peptides derived from the same CTA genes across different patients, but the exact same CTA peptides could be found in multiple patients (Suppl. Figure S5C).

Investigating the immunopeptidome in this cross-entity cohort therefore resulted in the discovery of a number of potential tumor-associated antigen candidates for immunotherapy.

### The majority of MS-based neoantigen candidates is derived from RNA sources

For the identification of neoantigen candidates, we have optimized our bioinformatics pipeline (6) by including novel tools such as an expanded mutation calling algorithm (34) and an improved mutation to peptide converter. The peptide identification algorithm pFind (23) was used with subsequent rescoring by the machine learning algorithm Prosit (21) (Figure 1). Neoantigen candidates had to pass our comprehensive post-processing pipeline, which is described in detail in the method section. By utilizing a Prosit-based rescoring workflow for our proteogenomic data, we could increase the total number of identified neoantigen candidates by 14 (Figure 5A).

**Figure 5.**
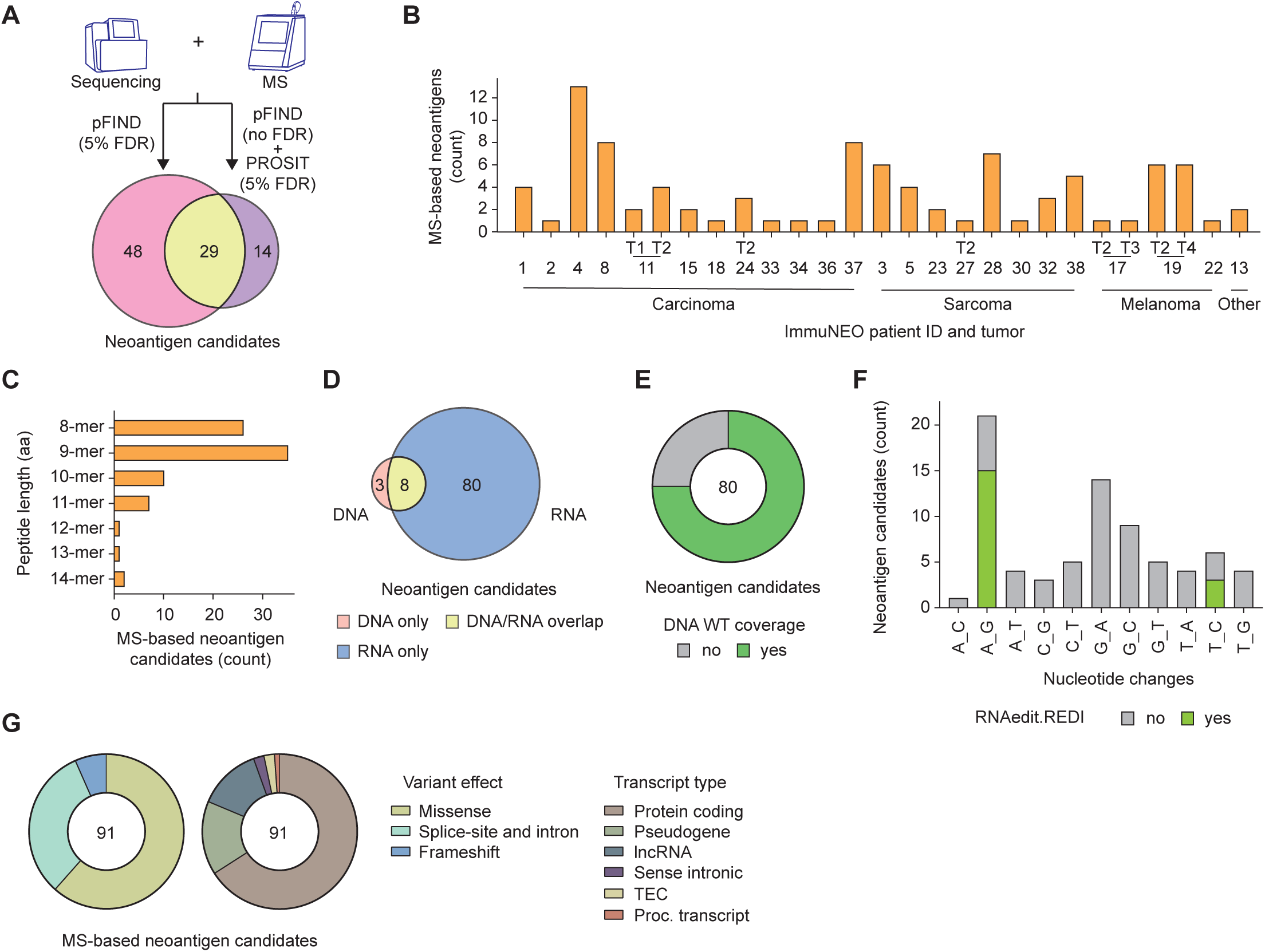
Proteogenomic identification of neoantigen candidates. **A, B,** Number of identified neoantigen candidates based on the bioinformatics tool that they were identified with (**A**) and per tumor sample and grouped by tumor entity (**B**). pFIND (v3.1.5) (23) was used at 5% FDR on spectral level for the identification of non-wild type (WT) 8-15mer neoantigen candidates. The machine learning tool Prosit (21) was additionally integrated to rescore the peptide spectra matching to the patient-specific ORF database using unfiltered pFIND data as input. n = 39 tumor samples from n = 32 patients were analysed in total; n = 27 tumor samples from n = 24 patients harboured n = 91 neoantigen candidates. **C,** Bar graph showing the length distribution of all identified neoantigen candidates in amino acids (aa). **D,** Genetic origin (DNA or RNA data) of the variants that the identified neoantigen candidates were derived from. **E,** Pie chart depicting the proportion of neoantigen candidates identified only from RNA sequencing (RNA-seq) data where the respective WT sequence was identified at the DNA level with a coverage of ≥ 3 reads (green) or the respective region was not covered at the DNA level (grey, < 3 reads). **F,** Distribution of the nucleotide exchange pattern of all variants that yield neoantigen candidates identified only from RNA-seq data. Variants previously identified in the REDIportal (29) database as RNA editing events are highlighted in green. **G,** Distribution of each mutation type (left) and biotype (right) of all variants that yield neoantigen candidates. **A-G,** n = 39 tumor samples from n = 32 patients were analysed in total; n = 27 tumor samples from n = 24 patients harboured n = 91 neoantigen candidates; n = 3 neoantigen candidates from DNA variants; n = 8 neoantigen candidates from DNA and RNA variants; n = 80 neoantigen candidates from RNA variants. aa, amino acids; MS, mass spectrometry; Proc., processed; T, tumor; TEC, to be experimentally confirmed; WT, wild type.

With this proteogenomic pipeline we were able to identify 91 neoantigen candidates in 24 patients across different tumor entities (75% of all patients) with 1 to 13 identified neoantigen candidates per patient (Figure 5B, Suppl. Table S4A), highlighting that most cancer patients harbor potential targets for personalized immunotherapy. We did not observe shared neoantigen candidates between patients, however, three peptides were shared between two metastases of a melanoma patient (ImmuNEO-19) and one peptide was shared between two distinct tumor samples of a patient with dMMR (ImmuNEO-11) (Suppl. Table S4A). Interestingly, we identified two neoantigen candidates in two patients (ImmuNEO-4 and -23) that were derived from shared genetic variants in MAP4K5 (IN_04_F, 1.5% FDR; shared between 32 tumor samples; Suppl. Table S2B) and in AC024075.2 (IN_23_A, 4.3% FDR, shared between 24 tumor samples; Suppl. Table S2B), respectively. Since both of these shared genetic variants were able to yield a pHLA-I that was presented in at least one patient, it is possible that these two peptides are presented in other patients with the genetic variants but were missed due to detection limitations of the patientś immunopeptidomes.

The peptide length of all identified neoantigen candidates ranged from 8 to 14 amino acids with nonamers predominating (Figure 5C). Perhaps most strikingly, out of 91 identified neoantigen candidates 80 were derived exclusively from RNA variants, while only three originated exclusively from DNA variants, and eight were shared between both sources (Figure 5D). Comparable to the overall number of RNA only variants, we could detect a corresponding WT sequence at the DNA level for the majority of identified neoantigen candidates that were derived exclusively from RNA variants (Figure 5E). Moreover, many of these variants also harbored an A to G nucleotide exchange pattern that has been associated with RNA editing and were reported as RNA editing events in the databank REDIportal (29) (Figure 5F). This suggests that RNA altering mechanisms (e.g. RNA editing) could be an important source for the formation of neoantigens. Regarding the variant effect of the variants that gave rise to the neoantigen candidates, missense variants were still most abundant, however, splice-site and intron variants were more prevalent compared to overall detected variants (Figure 5G, left). The majority of neoantigen candidates were derived from protein coding regions but a substantial amount was also derived from non-coding regions such as pseudogenes and lncRNAs (Figure 5G, right).

Taken together, our data indicate that MS-based identification of neoantigen candidates is feasible in the majority of cancer patients with tumor RNA representing an important source for the detection of peptide ligands derived from genetic variants.

### Identified neoantigens derived from RNA sources are immunogenic in a set of patients independent of the tumor entity

To assess the immunogenicity of the identified neoantigen candidates, we evaluated T cell responses against 79 neoantigen candidates from 21 patients in an *in vitro* assay with autologous or allogenic HLA-matched peripheral blood mononuclear cells (PBMCs) or expanded TILs by ELIspot analysis (Suppl. Figure S8A).

Out of 79 examined neoantigen candidates, 24 were capable of inducing T cell responses (29% of all tested neoantigen candidates) in either an autologous PBMC (Figure 6A, left), expanded TIL (Figure 6A, right), or an allogenic-matched PBMC (Figure 6B) culture setting (Figure 6C, Suppl. Table S4B). The majority of immunogenic neoantigens were identified by using autologous PBMCs and only three immunogenic neoantigens could be identified with expanded TILs (Figure 6A). This highlights the difficulties known for TIL cultures that could be explained by either insufficient expansion or a dysregulated and exhausted T cell phenotype of the expanded TILs, thus, preventing a proper T cell response against the presented neoantigen candidates. Although allogenic-matched PBMC cultures are challenging, especially with respect to donor selection, we tested a small set of neoantigen candidates (n=10) and could confirm the immunogenicity for four neoantigens that were immunogenic in the autologous setting and even identified one additional immunogenic neoantigen (IN_19_A) (Figure 6B). Of note, there was no enrichment observed regarding the frequency of immunogenic neoantigens out of the pool of neoantigen candidates that were identified by either of the two processing workflows or by both of them (Figure 5A, Suppl. Table S4B).

**Figure 6.**
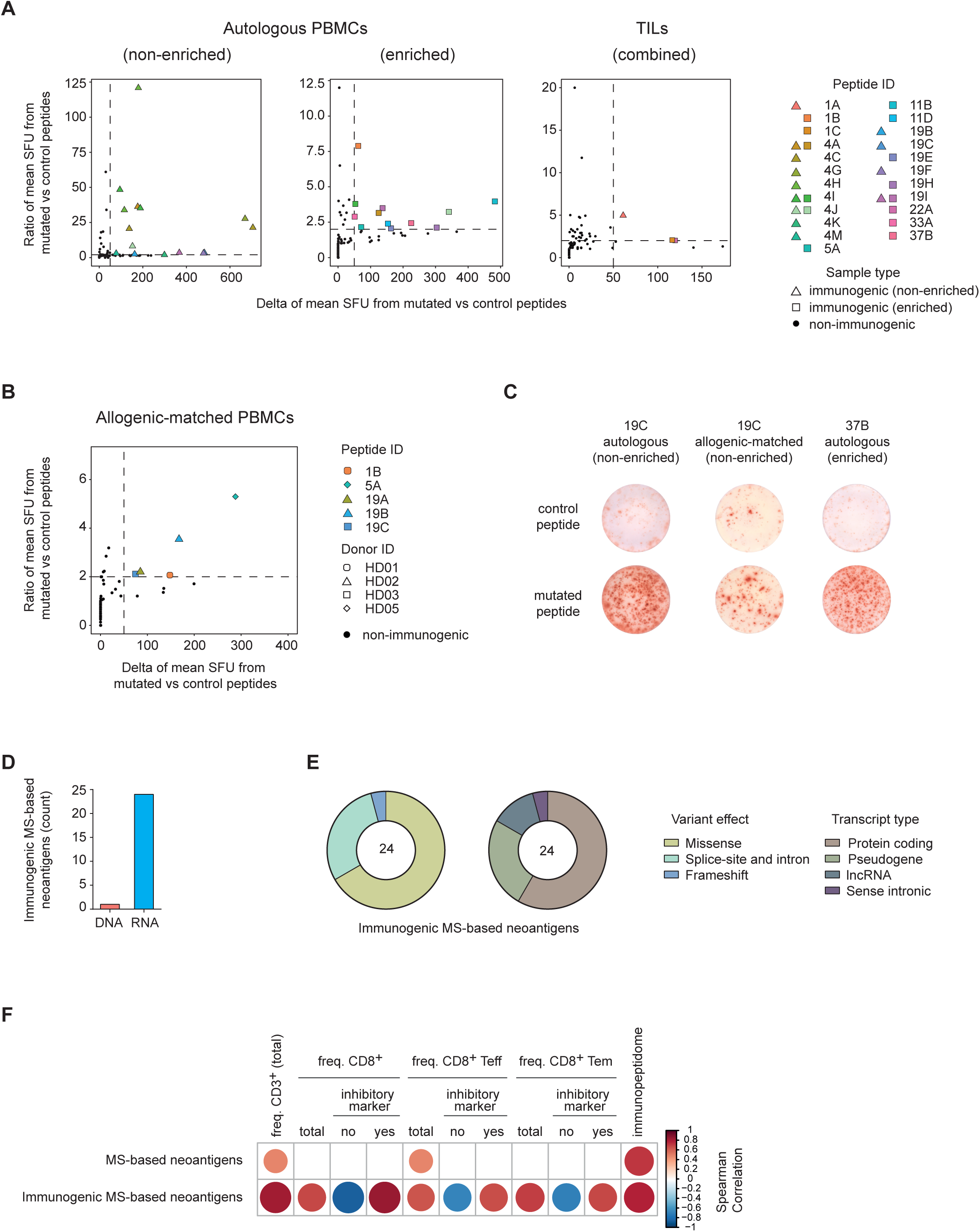
Immunogenicity assessment of neoantigen candidates A, B, Summary of immunogenicity assessment data from all performed modified accelerated co-cultured dendritic cell (acDC) assays for neoantigen candidates by ELIspot analysis using patient derived PBMC (non-enriched – left plot, CD137^+^ enriched – middle plot) or TILs (enriched and non-enriched combined – right plot) (**A**) and allogenic-matched healthy donor PBMCs (non-enriched) (**B**). Mean IFN-γ spot forming units (SFU) for T cells tested against the mutated peptide (test condition) and tested against a control peptide (control condition) were calculated and the ratio as well as the difference of the mean SFU have been determined. Values are shown for every peptide and PBMC or TIL aliquot tested. Highlighted are peptides that elicit an immune response where the ratio of SFU is > 2 and the difference of SFU is > 50. Autologous LCLs or allogenic HLA-matched cells (LCLs or HLA- transduced cell lines) were used as target cells. Negative values (when controls show more spots than the test condition) were set to 0 for better readability. **C,** Representative IFN-γ ELIspot data showing spots per well for autologous and allogenic-matched PBMCs tested against a control peptide (top) and the indicated neoantigen candidate (bottom). **D,** Genetic origin (DNA or RNA data) of the variants that the identified immunogenic neoantigens were derived from. **E,** Distribution of each mutation type (left) and biotype (right) of all variants that yield immunogenic neoantigens. **F,** Correlation matrix summarizing significant (p ≤ 0.05) Spearman correlations for multiple phenotypic parameters and the size of the immunopeptidome with the number of identified MS-based neoantigens overall and immunogenic ones. Spearman correlation coefficient Rho is labeled in color and size. For statistical analysis only one representative tumor sample per patient was used. **A, D-F,** n = 79 neoantigen candidates from n = 24 patients were analysed in total; n = 8 patients harboured n = 23 immunogenic neoantigens; n = 22 immunogenic neoantigen candidates from autologous PBMC cultures; n = 3 immunogenic neoantigen candidates from TIL cultures; n = 23 tumor samples from n = 17 patients for immunophenotyping data. **B,** n = 10 neoantigen candidates from n = 4 patients were analysed in total; n = 5 immunogenic neoantigen candidates from allogenic-matched PBMC cultures. MS, mass spectrometry; PBMCs, peripheral blood mononuclear cells; SFU, spot forming units; TIL, tumor- infiltration lymphocytes.

Importantly, all 24 immunogenic neoantigens were identified from RNA sources, with 23 detected exclusively from RNA variants and only one from both RNA and DNA variants (Figure 6D). In line with our findings for RNA only variants and neoantigen candidates, we observed that the majority of immunogenic neoantigens harbored a detectable WT sequence at the DNA level (Suppl. Figure S8B) and a substantial portion were reported as RNA editing events in the databank REDIportal (29) (Suppl. Figure S8C). This supports our hypothesis that RNA-altering mechanisms might be implicated in the formation of neoantigens that are capable of inducing T cell responses in patients. Moreover, the variant effect and the transcript type of the variants that gave rise to the immunogenic neoantigens were highly comparable to the distribution of neoantigen candidates as well (Figure 6E). When looking at binding predictions for our identified immunogenic neoantigens with NetMHC4.0 (35) and MHCFlurry (36) (Suppl. Table S4A), only 65% were identified as binders by at least one algorithm (percentile rank <2% or predicted binding affinity <500nM), indicating that one third would have been missed with these binding prediction algorithms. Overall, we observed immunogenicity of neoantigens regardless of the patients’ tumor entity, including patients with carcinoma, sarcoma, and melanoma (Suppl. Figure S8D, Suppl. Table S4B), indicating that the identification of immunogenic neoantigens is not limited to specific tumor entities.

Finally, in an effort to link the level of identified neoantigens (Figure 5 and 6) with the immune activity in the TME and the level of detected immunopeptides of our patients, we performed a Spearman’s rank correlation test with our immunophenotyping (Figure 2) and immunopeptidomic data (Figure 4). Since all neoantigen candidates were matched to the presence of pHLA-I mass spectra, both the number of neoantigen candidates and immunogenic neoantigens correlated strongly with the size of the detected immunopeptidome (Figure 6F). The overall number of neoantigen candidates also correlated slightly with the total frequency of CD3^+^ T cells and CD8^+^ Teff cells in the TME (Figure 6F). Importantly, the number of immunogenic neoantigens did not only correlate stronger with the total frequency of CD3^+^ T cells and CD8^+^ Teff cells in the TME, but we also observed a strong correlation with CD8^+^ T cells and CD8^+^ Tem cells as well as a generally more exhausted phenotype of these cells (Figure 6F). Thus, immunogenic neoantigens correlated with a more immune-active TME with high T-cell infiltration in our cohort.

In summary, we identified immunogenic neoantigens in a quarter of all patients of our pan-cancer cohort independent of the tumor entity by using a proteogenomic pipeline that utilizes RNA transcriptomics of tumor specimens for the identification of genetic variants. These immunogenic altered peptides correlated with T-cell infiltration and potentially an exhausted T cell phenotype.

## Discussion

The clinical application of personalized cancer immunotherapies based on neoantigens is benefitting greatly from the recent advances in mRNA-based vaccines (4) and cellular immunotherapy (37). However, the identification of tumor-specific and therapy-relevant targets is still critical. This is an area of research that mainly focused on cancer genomics and bioinformatics epitope prediction models for the identification of potential neoantigens in the past (1) but might benefit greatly from combinatorial approaches like proteogenomics that have been applied by other groups (7, 10) and us (6, 20). In this study, we showed that RNA is an important source for the identification of neoantigens and shared tumor antigens with our improved proteogenomic pipeline in an extensively characterized pan-cancer cohort. By combining proteogenomics with phenotypic and functional analyses, we linked the identified candidates to immunological features and validated their potential to induce T cell-driven immune responses.

Despite the relatively small size of this cohort and the high diversity with respect to tumor entity, disease stage, treatment history, age, and gender, we were able to confirm biomarkers with prognostic significance which have been already established for a number of distinct malignancies, indicating that these biomarkers have a strong prognostic power. When looking at the TMB as a prognostic biomarker, we could confirm a significant positive correlation between the number of somatic mutations and patients’ survival, as previously shown for several different cancer entities as well as selected cross-entity studies (38–41). In addition, we observed that high levels of CD8^+^ T cells expressing inhibitory markers, previously shown as an indication for a dysfunctional T cell state in the TME (42), correlated with poor clinical outcome.

To increase the number of identified neoantigens from our previously published proteogenomic strategy (6), we integrated tumor RNA as an additional source for variant detection. Including RNA-seq to our pipeline has two advantages. First, RNA-seq has been shown to complement WES in calling somatic mutations in glioblastoma multiforme to broaden the scope of discoveries (43). Second, RNA-seq is able to detect variants that are not occurring at the DNA level but are derived from RNA processing events like alternative splicing and RNA editing (44, 45). It has been previously reported that RNA editing events and RNA dysregulation lead to the diversification of the cancer proteome (14, 15) and in fact, we substantially increased the number of genetic variants and neoantigens by including RNA-seq in our pipeline. Variant detection using RNA-seq is already utilized in a number of studies for the identification of neoepitopes (7, 16) but comes with its own limitations, in particular for variants derived from RNA processing events since they cannot be validated by matched-normal DNA samples. In addition, obtaining matched-normal RNA samples from the same tissue as the tumor is similarly limited as it might be either not available or may be influenced by the tumor activity and transcriptional profile of the surrounding tissue. To exclude false positive RNA variants based on single-nucleotide polymorphisms (SNPs), we used a methodology of combining tumor RNA-seq with normal WES data that has been shown to be most effective for calling RNA variants (46). We thereby excluded frequent population SNPs. Since that still did not control for false positive RNA variants from RNA processing events, we overcame this limitation by matching the RNA variants to the MS spectra from the tumor pHLA-I and thereby performed a cross-validation of the neoantigen candidates. Of note, due to this subsequent cross-validation, less stringent mutation calling algorithms for RNA but also DNA variant detection were used that increase the search space for potential neoantigen identification. Therefore, false positive hits may have been still not completely excluded here. However, using our sensitive algorithms for the detection of genetic variants, we were able to identify genetic variants that occurred not only in individual patients but were shared in a substantial number of patients at the DNA and especially at the RNA level, representing potentially attractive common targets that need to be investigated in a larger cross-entity cohort.

The strength of our neoantigen discovery platform, the matching of MS-spectra to variants, is also its bottleneck because the number of identified neoantigens strongly correlated with the size of the immunopeptidome. Therefore, improving MS-based neoantigen detection is paramount and there are three avenues that can be addressed. (1) Optimizing artificial intelligence tools for the matching and rescoring of MS spectra (like Prosit) will enhance their potential for neoantigen discovery. (2) Improving protocols for sample preparation and immunoprecipitation of pHLA-I might result in a higher yield of detected peptides. (3) Increasing the sensitivity of MS instruments will likely have the biggest impact in the future (47).

The number of neoantigen candidates that we identified was small compared to the thousands of hits that were reported with epitope prediction models (1, 2). However, in our study approximately 30% of the tested candidates elicited a T cell response *in vitro*, a far greater number than could be expected from any epitope prediction approach. Thus, drastically reducing the need for large-scale immunogenicity testing that would not be feasible in a clinical environment. Since a substantial portion of our immunogenic neoantigens was not predicted as binders, solely prediction-based approaches might miss these potentially promising targets. Moreover, immunogenicity testing in autologous T cell assays has the inherent risk of a lack of an immune response to the presented peptide because of T cell dysfunction (48), suggesting that some neoantigen candidates that did not induce a T cell response might actually be potentially immunogenic. More sensitive assays for validation are therefore necessary and combined single cell RNA and T cell receptor (TCR) sequencing shows great promise for this need (48, 49).

Neoantigen candidates derived from RNA variants have been previously reported (7–10,16,17) and may represent missing targets in studies where suspected neoantigens could not be detected by focusing only on WES (49). Indeed, the majority of neoantigens in our cohort were derived only from RNA variants and we observed a high number of A to G modifications typical for RNA editing events (16, 28). However, RNA variants detected in this cohort also include other forms of RNA dysregulation as well as potentially somatic mutations which have not been covered by WES. Elucidating the nature of RNA variants and their role in cancer biology and immunotherapy is an important research area (reviewed in (50–52)) that might lead to new types of cancer treatment.

Neoantigen-based vaccines showed limited clinical response in previous trials (5, 53). This might have been due to poor candidate selection or because of a dysregulated T cell state in the treated patients. However, some efficacy has recently been observed using mRNA vaccination in melanoma and pancreatic cancer including also a combination with immune checkpoint inhibitors (4,54,55), suggesting that it is crucial to overcome the dysregulated T cell state for neoantigen vaccines to be efficacious. It will be important to understand subtle differences in vaccines and clinical protocols in order to understand outcomes of these early trials. In addition, developing alternative strategies that engage non-dysfunctional T cells like neoantigen-specific TCR-T cell therapy is of great importance to treat patients that do not respond to immune checkpoint inhibition.

Taken together, our data identified a number of attractive cancer-associated and -specific canonical and non-canonical peptide antigens that have been partially shared by a significant portion of patients in our cohort. Most importantly, we demonstrate the importance of RNA as a source for MS-based neoantigen identification in a large number of patients of this cross-disease cohort correlating with T-cell infiltration. Functionally active neoantigen-specific T cells could be identified only in a sub-cohort of these patients likely due to a severe dysfunctional state of these T cells. Therefore, immunotherapies focusing on the rescue of such T cells or targeting neoantigens with a non-dysfunctional repertoire including TCR-transgenic T cells may represent a valid immunotherapeutic option for a large number of cancer patients.

## Methods

### Primary human material and cell lines

Informed consent of all participants was obtained following requirements of the institutional review boards (Ethics Commission of the Medical Faculty of Technical University Munich and Ethics Committee of the Medical Faculty of Heidelberg University (S-206/2011)). An overview about all patients is given in Supplementary Table S1A. Tumour tissue samples were collected from patients, who underwent tumor resection at the different DKTK partner sites. Immediately after resection, fresh tumor tissue was macroscopically dissected by an experienced pathologist and stored in PBS at 4°C for transport or until processing. Additional tumour tissue was formalin-fixed and paraffin-embedded (FFPE). Before molecular analysis, tumor diagnosis was confirmed by a pathologist and tumor content was determined by an HE stain taken from the sample going to be used.

From the fresh tumour tissue a part was snap frozen and stored in liquid nitrogen (-196 °C) for later sequencing and mass spectrometry analysis.

From all remaining fresh tissue a single cell suspension was generated by mincing and digesting 0.2g tissue pieces per tube for 90min at 37°C in 1ml RPMI supplemented with 40µL Enzyme H (Tumor dissociation kit human, Miltenyi; Stock conc.), 5µL Enzyme A (Tumor dissociation kit human, Miltenyi; Stock conc.), 25µL Hyaluronidase (Sigma Aldrich, 10 mg/mL stock), 25µL DNAse I (Sigma Aldrich, 10 mg/mL stock). After digest the suspension and tissue pieces were meshed, and single cells were used for flow-cytometry analysis and FACS analysis.

Primary patient cells used in this study: For TIL generation, part of the fresh tumor tissues was minced and TILs were expanded for 2–3 weeks by cultivation with irradiated feeder PBMC, 1000 U/ml IL-2 (PeproTech) and 30 ng/mL OKT3 (kindly provided by Elisabeth Kremmer). Change of medium supplemented with 300 U/mL IL-2 was performed twice a week. After expansion for 2 weeks, TILs were frozen for later use in stimulation assays. PBMC from patients were isolated from whole blood by density-gradient centrifugation (Ficoll/Hypaque, Biochrom) immediately on receipt and frozen for later use in stimulation assays. Patients’ T cells, derived from PBMCs or TILs, were cultivated in T-cell medium (TCM): RPMI 1640 (Invitrogen) supplemented with Penicillin/Streptomycin (Pen/Strep) (Invitrogen), 5% FCS (Invitrogen), 5% human serum (HS), 10 mM Hepes (Invitrogen), 10 mM MEM non-essential amino acids (Invitrogen), 1 mM MEM sodium-pyruvate (Invitrogen), 2 mM L-Glutamine (Invitrogen) and 16.6 µg/mL Gentamycin (Biochrom).

Cell lines used in this study: T2 and C1R cell lines (American Type Culture Collection (ATCC) and lymphoblastoid cell lines (LCL) generated from patient samples (LCL IN-01, IN-03, IN-04, IN-08, IN-09, IN-11, IN-13, IN-18, IN-19, IN-22, IN-24, IN-33, IN-37) and healthy donors (HD) (LCL HD04, HD06, HD07, HD08) or purchased from ATCC (LCL CLA, Daudi, FM, IBW9, RSH, SWEIG007) were used. Morphology and constant growth behaviour of all cell lines were controlled periodically, and the absence of mycoplasma infection was routinely confirmed by PCR (Venor GeM mycoplasma detection kit, Minerva Biolabs). T2 and C1R were retrovirally transduced with the HLA restriction elements HLA-A6601 (C1R- A6601), B0702 (C1R-B0702), A0301 (T2-A0301), B5101 (T2-B1501) and B4402 (T2-B4402) as described before (6). All target cell lines were maintained in complete RPMI (cRPMI): RPMI 1640 (Invitrogen) supplemented with Pen/Strep (Invitrogen), 10 mM MEM non-essential amino acids (Invitrogen), 1 mM MEM sodium-pyruvate (Invitrogen), 2 mM L-Glutamine (Invitrogen) and 10% FCS (Invitrogen).

### Whole exome and RNA sequencing of patient material and analysis

#### Extraction of nucleic acids

DNA and RNA from tumor specimens and DNA from matched blood samples were isolated using the AllPrep DNA/RNA/miRNA Universal Kit (Qiagen). For formalin-fixed and paraffin-embedded (FFPE) samples, the AllPrep DNA/RNA FFPE Kit (Qiagen) was used. DNA from blood samples was isolated using the QIAsymphony DSP DNA Mini Kit (Qiagen) or the QIAamp DNA Blood Mini Kit (Qiagen). Quality control and quantification were done using a FilterMax F3 Multi-Mode Microplate Reader (Molecular Devices), a 4200 or 2200 TapeStation system (Agilent).

#### Library preparation and target capture for whole-exome sequencing

For whole-exome sequencing (WES) library preparation, 1.5 µg genomic DNA were fragmented to 150- 200 base pair (bp; paired-end) insert size with a Covaris S2 device, and 250 ng of Illumina adapter-containing libraries were hybridized with exome baits at 65°C for 16 hours. Exome capturing was performed using SureSelect Human All Exon in-solution capture reagents (Agilent). In case RNA was pooled in for sequencing, V5 without UTRs was used to reach a minimum average coverage of 80x for the tumor and 50x for the control. V5 with UTRs was used when DNA was sequenced alone.

#### Library preparation for whole-genome sequencing

Whole-genome sequencing (WGS) libraries were prepared using the TrueSeq Nano Library Preparation Kit (Illumina) following the manufacturer’s instructions.

#### Library preparation for RNA sequencing

RNA sequencing (RNA-seq) libraries were prepared using the TruSeq RNA Sample Preparation Kit v2 (Illumina) using the stranded protocol. Briefly, mRNA was purified from 1 µg total RNA using oligo(dT) beads, poly(A)+ RNA was fragmented to 150 bp and converted into cDNA, and cDNA fragments were end-repaired, adenylated on the 3’ end, adapter-ligated, and amplified with 12 cycles of PCR. 2 The final libraries were validated using a Qubit 2.0 Fluorometer (Life Technologies) and a Bioanalyzer 2100 system (Agilent).

#### Whole-exome, whole-genome, and RNA sequencing

Paired-end sequencing (2 x 150 bp) was performed with HiSeq X-Ten instruments (Illumina). Two lanes, each of tumor and control, were sequenced, yielding an average coverage of at least 70x for WGS cases. Paired-end sequencing (2 x 100 bp) was carried out with HiSeq 4000 (Illumina), pooling two patients’ samples on one lane. From January 2017, RNA was sequenced separately with dual indexing in pools of three samples per HiSeq 4000 lane or multiplexed over several lanes to prevent adapter hopping. From October 2019, RNA was sequenced in pools of 3-5 samples per NovaSeq 6000 lane. Comparability of data has been validated.

### Mutation calling from exome and RNA sequencing data

Mutation calling was performed on WES/WGS and RNA-Seq data for identification of single nucleotide variants and insertion/deletions for the indicated patients (Suppl. Table S1A). Analysis of WES data was performed following the GATK Best Practice suggestions and utilizing the established analysis pipeline MoCaSeq (34), adapted for the human genome. After read trimming using Trimmomatic 0.38 (LEADING:25 TRAILING:25 MINLEN:50), bwa mem 0.7.17 was used to map reads to the human reference genome (GRCh38.p12). Picard 2.18.26 and GATK 4.1.0.0 were used for postprocessing (CleanSam, MarkDuplicates, BaseRecalibrator) using default settings. Somatic mutations were called using MuTect2 4.1.0.0 (56). SNVs and Indels ≤ 10 base pairs were annotated using SnpEff 4.3t, based on Ensembl 92.

For mutation calling from RNA-Seq, raw reads were trimmed using Trimmomatic (LEADING:25 TRAILING:25 SLIDINGWINDOW:10:25 MINLEN:50) and aligned to the human reference genome with STAR (2.6.0c). Mutations were called using Strelka2 (2.9.10) using the RNA option (57). SNVs and Indels ≤ 10 base pairs were annotated using SnpEff 4.3t, based on Ensembl 92. De novo variant calling on tumor WES data was performed by comparison to PBMC WES data.

For variant calling on RNA-Seq data, positions sufficiently covered in WES with no evidence for the presence of germline SNVs/indels, were included as somatic. Furthermore, for positions where SNVs/indels were called only by Mutect2 or Strelka2, the threshold to include this SNV/indel in the second tissue sample was substantially lowered and did not require to be called separately by Mutect2/Strelka2.

Population SNPs with certain population allele frequency based on GnomAD (58) (>1%) and dbSNP (>5%) (59) were excluded.

To calculate the tumor mutational burden (TMB), first WES probe regions (+/- 300bp) with a coverage above 10 reads were identified. The TMB was then calculated as the number of genic/non-synonymous mutations overlapping with these regions divided by the total length of probe regions in megabases (Mb).

### HLA typing

HLA typing was done from the available whole exome or whole genome sequencing data using the consensus of all xHLA (60), BWAKit (61) and OptiType (62) using default settings. For confirmation, HLA typing was done on gDNA isolated from PBMC by targeted next generation sequencing in selected patients (Zentrum für Humangenetik und Laboratoriumsdiagnostik, Martinsried, Germany).

### Immunoprecipitation of HLA complexes and liquid chromatography (LC)-MS/MS analysis of eluted peptides

Immunoprecipitation of HLA complexes, consequent elution and purification of peptide ligands was performed on indicated tumor samples (Suppl. Table S1A) as previously described (Bassani-Sternberg et al., 2016). Briefly, snap-frozen tumor tissue samples were placed in 5-7 ml of PBS with 0.25% sodium deoxycholate (Sigma-Aldrich), 1% octyl-β-D glucopyranoside (Sigma-Aldrich), 0.2 mM iodoacetamide, 1 mM EDTA, and 1:200 Protease Inhibitor Cocktail (Sigma-Aldrich) and mechanically dissociated with an ULTRA-TURRAX Disperser (IKA) for 10 s on ice, followed by 1 h incubation at 4°C. The lysates were then cleared by centrifugation at 40,000g at 4 °C for 20 min and flowed through columns packed with protein-A Sepharose beads (Invitrogen) to deplete the endogenous antibodies. HLA class I complexes were immunoaffinity-purified from the cleared and antibody-depleted lysates on columns containing protein-A Sepharose beads covalently bound to 2 mg of the pan-HLA class I antibody W6/32 (purified from HB95 cells; ATCC) and eluted at room temperature with 0.1 N acetic acid. The eluted HLA-I complexes were then loaded onto Sep-Pak tC18 cartridges (Waters Corporation), and HLA-I peptides were separated from the complexes by elution with 30% acetonitrile (ACN) in 0.1% trifluoroacetic acid (TFA). Peptides were further purified using Silica C-18 column tips (Harvard Apparatus), eluted again with 30% ACN in 0.1% TFA and concentrated by vacuum centrifugation. Finally, HLA-I peptides were resuspended with 2% ACN in 0.1% TFA for LC-MS/MS analysis.

LC-MS/MS analysis was performed on an EASY-nLC 1200 system (Thermo Fisher Scientific) coupled online with a nanoelectrospray source (Thermo Fisher Scientific) to a QExactive HF-X mass spectrometer (Thermo Fisher Scientific). Peptides were loaded in buffer A (0.1% formic acid) on a 50 cm long, 75 µm inner diameter column, in-house packed with ReproSil-Pur C18-AQ 1.9 µm resin (Dr. Maisch HPLC GmbH), and eluted during a 95 min linear gradient of 5-30% buffer B (80% ACN, 0.1% formic acid) at a flow rate of 300 nl/min. The mass spectrometer was operated in a data-dependent mode with the Xcalibur software (Thermo Scientific). Full MS scans were acquired at a resolution of 60,000 at 200 m/z and AGC target value of 3e6 with a maximum injection time of 80 ms. The ten most abundant ions with charge 1-4 were accumulated to an AGC target value of 1e5 and for a maximum injection time of 120 ms and fragmented by higher-energy collisional dissociation (HCD). MS/MS scans were acquired with a resolution of 15,000 at 200 m/z and 20 s dynamic exclusion to reduce repeated peptide selection.

### Wild-type peptidome analysis

For the identification of peptide sequences from the MS spectra, pFIND 3.1.5 (23) was used to match the reference protein database (Human Ensembl GRCh38, release 92) with general contaminants to the generated spectra files. Parameters were set to search for non-specifically digested peptides ranging from 8-15mers with a maximum mass of 1,500 Da and N-terminal protein acetylation (42.010565 Da), methionine oxidation (15.994915 Da), cysteine carbamidomethylation (57.021463 Da) as possible post-translational modifications. Identified peptides were filtered with an FDR of 0.01 (and 0.05) at the peptide spectrum match level.

### MHC-motif deconvolution

To assess the quality and purity of the MS-generated immunopeptidomic data, the identified peptide sequences where deconvoluted to the respective patients HLA-allele by their binding motif using MHCMotifDecon-1.0 (63, 64). Here, MHC binding predictions from NetMHCpan-4.1 (for MHC class I) are used to deconvolute and assign likely MHC restriction elements to MHC peptidome data. All identified peptide sequences with lengths of 8-15 amino acids and all HLA-A, B and C alleles of each patient have been used for analysis applying standard setting as indicated on the website.

### Pipeline for the identification of patient specific neoantigen candidates from MS data

In order to improve the identification of neoantigens we further developed our MS-based pipeline (6) for the analysis of this diverse patient cohort (Figure 1). The following novel features have been integrated: (1) On the genetic level, mutation calling from RNA sequencing data has been accomplished using Strelka2 (34). Moreover, a refined algorithm for translation of open reading frames (ORFs) in all three frames has been implemented to identify potential neoantigens from a large source of genetic aberrations (splice site variants, intron-inclusions, non-coding variants, etc.). (2) On the proteomic level, pFIND as a peptide calling tool (23) as well as the machine learning tool Prosit (20, 21) have been included into the pipeline. (3) We additionally established a comprehensive post-processing filtering procedure, especially focusing on exclusion of possible wt peptides and SNPs. In detail, the subsequently described analysis steps have been performed.

#### Generation of custom database for MS-based identification of mutated peptides

With the main goal to obtain mutated peptide sequences, mutations called from WES/WGS and RNAseq were introduced into the wildtype transcript DNA sequences downloaded from biomart (v92) and translated into peptide sequences. Genes were included in the analysis without exceptions regarding the transcript biotypes. For non-protein-coding transcripts, ORFs enclosing the mutation site were determined by identifying paired start and stop-codons in all 3 reading frames. The same procedure was performed for protein-coding transcripts in case of start/stop-loss/gain and frameshift mutations. Furthermore, for start/stop mutations, the coding sequence (CDS) was extended into the corresponding UTR. For mutations affecting splice donor or acceptor sites, the affected intron was included into the CDS and again checked for valid ORFs. Only mutations resulting in amino acid changes and within valid ORFs were considered. For every affected transcript, up to three ORFs enclosing the mutation site were translated into the corresponding mutated peptide sequence. Peptide sequences were then used together with the immunopeptidomics data from mass spectrometry.

#### Identification of mutated peptides sequences from MS data

For the identification of mutated HLA class I peptides, the reference protein database (Human Ensembl GRCh38, release 92) was searched together with the patient-specific customized databases containing the mutated sequences from step 1 using pFIND 3.1.5 (23). Parameters were set to search for non-specifically digested peptides ranging from 8-15mers with a maximum mass of 1,500 Da and N-terminal acetylation (42.010565 Da), methionine oxidation (15.994915 Da), cysteine carbamidomethylation (57.021463 Da) as possible post-translational modifications and a set FDR of 0.05 at the peptide spectrum match level. After protein annotation, the pFIND generated unfiltered peptide lists were (1) filtered for the FDR of 0.05 and used directly for further post-processing (pFIND peptides) and (2) used unfiltered for subsequent re-scoring and analysis by the Prosit pipeline (Prosit peptides) (20, 21). The rescoring method is extensively described in Wilhelm *et al.*, 2021. In brief, the unfiltered search engine output including decoys of pFind was used as input for the spectral intensity-based rescoring. Unprocessed MS2 spectra corresponding to the identifications were annotated with all matching b- and y-ions. Spectral comparison between predicted fragment ion intensities and experimental intensities was performed for using the best-matching prediction settings and calculating previously described similarity measures (e.g., normalized spectral contrast angle). FDR estimation was performed using SVM Percolator 3.00 (65). All PSMs surpassing a FDR threshold of 5% were further considered for analysis.

Peptides identified by both approaches were combined and used for post-processing.

### Post-processing and filtering of neoantigen candidates and MHC binding prediction

Peptides were filtered to remove contaminants and reverse sequences, which were only used to determine statistical cutoffs. In addition, the results were filtered for sequences identified exclusively in the custom mutated databases, and not in the Ensembl database, thus ensuring the peptide originating from a non-wild-type ORF.

Peptides harboring mutations (SNVs, In/Dels, multiple substitutions) within their sequence were directly taken as valid, whereas peptides not containing the mutations in the peptide sequence were further assessed. SNVs outside of the peptide sequence were excluded, whereas frameshift mutations upstream of a peptide or splice site mutations were checked manually in BLAT (66) and were considered “mutated” or “non-wt” if a peptide within a noncanonical frame or a retained intron was detected. The filtered potential neoantigens were then checked via an automated protein BLAST (67) search and peptides with more than 2 hits in the protein data base were excluded while peptides with 1-2 hits were double checked manually by literature research and excluded if necessary. Additionally, three different peptide data bases PeptideAtlas (68), PepBank (69) and IEDB (70) were used to filter for already known (immunogenic) peptides.

After complete filtering the binding affinity of each neoantigen candidate was predicted by using two different algorithms, NetMHC 4.0 (35) and MHCflurry 1.6.0 (models class1) (36), and the best binding allele according to predicted affinity or percentile rank was determined for each algorithm.

### Flow cytometry analysis of tumor single cells and FACS sort

For flow cytometry analysis, up to 0.5 Mio alive single cells from the digested tumor tissue have been used per panel and isotype controls. Cells were first incubated in 50µL human serum (HS) for 20min for blocking unspecific binding. Subsequently, ethidium monoazide bromide (EMA, 1:500, Thermo Fisher Scientific) was added for live-dead staining to the HS and incubated 10min on ice in the dark and 10min on ice in the light. After washing 2µL of the respective antibodies or 1.5µL of the isotype control antibodies were added and stained for 20min on ice in the dark. The following antibodies were used: CD45-PerCP-Cy5.5 (clone HI30, ref. 564105, BD), CD3-AF700 (clone UCHT1, ref. 300423, BioLegend), CD8-APCH7 (clone SK1, ref. 560179, BD), CD4-V450 (clone SK3, ref. 651849, BD), CD45RA- BV510 (clone HI100, ref. 304141, BD), CD62L-PE (clone DREG-56, ref. 560966, BD), CD366-BB515 (anti-TIM-3, clone 7D3, ref. 565568, BD), CD279-PECy7 (anti-PD-1, clone EH12.2H, ref. 329917, BioLegend), CD223-APC (anti-LAG-3, clone 3DS223H, ref. 17-2239-42, eBioscience), CD103-FITC (clone Ber-ACT8, ref. 550259, BD), HLA-DR-APC (clone G46-6, ref. 559866, BD), CD56-PE (clone 5.1H11, ref. 362508, BioLegend), CD45-APC-H7 (clone 2D1, ref. 560274, BD), CD25-PE (clone 2A3, ref. 341011, BD), CD127-BV510 (clone A019D5, ref. 351331, BioLegend). Appropriate isotype controls for each antibody were used as negative control. After staining cells were washed and fixed with paraformaldehyde (PFA, 1%, Sigma Aldrich) and stored at 4°C for later analysis. Measurements were performed on an LSR II (BD) and anti-IgG beads (Miltenyi) as well as unstained cells were used for single stains and instrument set- up. Voltages were adapted to the autofluorescence of each patient tumor and all possible events were measured using FACS DIVA software. All steps were carried out on ice and as quickly as possible to minimize changes in cell viability and marker expression. Data analysis and compensation was performed using FlowJo V10.7.1 and the gating strategy was kept consistent for every sample depending on the panels analysed (Gating strategy see Suppl. Figure S2A).

For sorting of CD8^+^ T lymphocytes, min. 5-10 Mio cells were taken from the digested tumor sample/single cell suspension (when enough cells were available). Cells were blocked with 200-500µL HS depending on the cell numbers for 20min on ice in the dark. After washing 2µL/1Mio cells of the respective antibodies, CD8-PECy7 (clone RPAT-8, ref. 557746, BD) and CD45-APC (clone J33, ref. IM2473, Beckman Coulter), and 7-Amino-Actinomycin D (7AAD, Invitrogen) for live-dead staining were incubated in 100-200µL FACS Buffer for 30min on ice in the dark. After washing cells were resuspended in 1mL/10Mio cells FACS buffer, filtered and directly used for sorting on a FACSAria III (BD). Single stains were generated using anti-IgG micro beads (Miltenyi) according to the manufacturer’s instructions and were used together with unstained cells and 7AAD-only stained cells for on-device compensation. Alive-SingleCells-CD45^+^-CD8^+^ cells were sorted into pre-cooled tubes containing RPMI. Sorted cells have been pelleted and resuspended in 300µL RNA Protect (Qiagen), snap frozen and stored in liquid nitrogen (-196 °C) for later mRNA sequencing analysis. All steps have been carried out on ice and as quickly as possible to minimize changes in cell viability and marker expression.

### Bulk mRNA sequencing of sorted cells and analysis

Paired-end sequencing (2 x 75bp) was performed on a NextSeq 500 (Illumina) with SMART-Seq Stranded Kits (Takara Bio, USA) to reach at least 50 Mio. raw reads per sample. The raw sequencing data was processed with Trimmomatic version 0.36 (71). Trimmed reads were acquired by removing Illumina TruSeq3 adapters and bases at the start and end of each read, for which the phread score was below 25. Further reads were clipped if the average quality within a sliding window of 10 fell below a phread score of 25. Conclusively reads smaller than 50 bases were removed. For mapping and counting, the human gene annotation release 29 and the corresponding genome (GRCh38.p12) were derived from the GENCODE homepage (72). STAR version 2.7.5b (73) was used to map the trimmed sequencing data to the reference genome, with the parameters adapted from protocol recommendations (74). Mapped reads were deduplicated with bamUtils v1.0.14 (75) and featureCounts v.1.6.3 (76) was used to assign and summarize reads to genes while ignoring multi-mapping, multi-overlapping and duplicated reads. The resulting raw count matrix was imported into R v4.0.5 and lowly expressed genes were subsequently filtered out. Prior differential expression analysis with DESeq2 v1.18.1 (77), dispersion of the data was estimated with a parametric fit using the Survival as explanatory variable. Shrunken log2 fold changes were calculated afterwards with the apeglm method (78) and used as ranking criteria for the pathway analysis with GSEA in the preRanked mode (79). The Hallmark and Gene Ontology gene set definitions from MsigDB v7.4 (80, 81) were used for GSEA.

### Generation of lymphoblastoid cell lines as autologous target cells

For the generation of patient derived LCL, first potent Epstein-Barr virus (EBV) supernatant was generated from B95-8 cells (provided by Ulrike Protzer). Therefore, 1 Mio cells per mL were stimulated in cRPMI (see Methods – primary human material and cell lines) with 20 ng/mL PMA (Sigma Aldrich) for 1h at 37°C, subsequently washed 3 times and cultured at a concentration of 1 Mio cells per mL in fresh cRPMI. After 3 days the supernatant was harvested, filtered with a 0.45µm sterile filter and stored at -80°C for up to 1 year. Afterwards, this supernatant was used for the infection and immortalization of patient derived B cells from PBMC samples. Therefore, up to 0.5 Mio PBMCs were incubated in 1mL RPMI with 1mL EBV supernatant for 2h at 37°C, following the addition of further 1mL cRPMI supplemented with Cyclosporine A (Sigma Aldrich) to a final concentration of 1µg/mL and culture in cell culture flasks at 37°C. Cells were split once clusters were visible and/or medium colour changed and expanded at 0.3-0.6 Mio cells per mL until enough cells were available for freezing or direct use in experiments.

### Immunogenicity assessment of identified peptide ligands

Recall antigen-experienced T cell-responses to selected peptides were investigated as previously described with modifications (6, 82). In brief, up to 1 Mio PBMCs or TILs per well from each patient were used for in vitro screening. For peptide stimulation on day 0, 1 µM of each synthetic peptide (>90% purity, DGPeptidesCo Ltd.) was added to the culture along with 0.5 ng/ml Interleukin (IL)-7 (Peprotech), 50 ng/ml Tumor necrosis factor (TNF)-α (Peprotech) and 10 ng/ml IL-1β (Peprotech). As positive control T cells have been non-specifically stimulated with 0.5 ng/µL phorbol-12-myristate-13- acetate (PMA, Sigma Aldrich) and 1 ng/µL Ionomycin (Sigma Aldrich). After 24h of peptide stimulation, 100µL supernatant was collected for later ELISA analysis and cells where either used for direct overnight ELISpot analysis as previously published or enriched for specifically activated T cells using a CD137^+^-based magnetic isolation (83). CD137-expressing activated cells were isolated and enriched using the human CD137 MicroBead Kit (Miltenyi) according to the manufacturer’s instructions. Enriched cells were taken into culture in T cell medium (TCM, see Methods – primary human material and cell lines) supplemented with 5 ng/mL IL-7 (Peprotech), 5 ng/mL IL-15 (Peprotech), 30 U/mL IL-2 (Peprotech) and 30 ng/mL OKT-3 (kindly provided by Elisabeth Kremmer) along with 1 Mio irradiated (30 Gray) feeder PBMC. Enriched cells were cultured for 12 days and fed by adding IL-7 and IL-15 twice per week and IL-2 once per week. Non-enriched cells were cultured and expanded in TCM supplemented with 5 ng/mL IL-7 (Peprotech) and 5 ng/mL IL-15 (Peprotech) and fed twice per week. For assays using healthy donor PBMCs the protocol without enrichment was followed and a different HLA-matched donor for each peptide was selected based on the affinity predictions performed by NetMHC 4.0 (35) and MHCflurry (36), where possible.

After 13 days of expansion, reactivities of T cells to the synthetic peptide ligands was assessed by specific interferon (IFN)-γ release by ELISpot assay. As antigen-presenting target cells for the second stimulation on day 13, either an autologous lymphoblastoid cell line (LCL) derived from the same patient or HLA-matched LCL, HLA-transduced T2 or C1R cells were used. The target cells were pulsed for 2 h with either the selected mutated peptide or an control peptide prior to co-culture with the T cells (in duplicate or triplicate according to available cell numbers). The co-cultures were performed with an effecter-to-target ratio of 2:1 using 20,000 pre-stimulated T cells and 10,000 pulsed target cells per well. ELISpot plates (Merck Millipore) were coated with an IFN-γ capture antibody (1-D1K, Mabtech) at 4°C overnight prior to the co-culture, development was performed with an IFN-γ- detection antibody (7-B6-1-biotin, Mabtech) and Streptavidin-HRP (Mabtech). ELISpot plates were read out on an ImmunoSpot S6 Ultra-V Analyzer using Immunospot software 5.4.0.1 (CTL-Europe).

We defined the reactivity by the spot counts at day 13 comparing the mean spots from the mutated peptide condition with the mean spots from the control peptide condition and set the threshold to a ratio above 2, meaning the mutated peptides elicited an IFN-𝛾 response in twice as many T cells compared to the control, and a difference of spots above 50, which is defined as the background threshold.

### Statistical analysis

Two-tailed Mann-Whitney U test was used to compare frequencies of CD8^+^ T cells expressing at least one activation marker (HLA-DR, CD103) of tumor specimens with high vs low immune cell infiltration.

Correlations of two distinct parameters were assessed using Spearman’s rank correlation coefficient. For correlation of the numbers of DNA variants and RNA variants, all samples with both analyses were included while for the correlation of phenotypic data with peptidomic data only one representative tumor sample from each patient included for analysis (ImmuNEO core cohort see Suppl. Table S1A) has been used to circumvent bias due to multiple metastasis available for some patients.

Kaplan-Meier curves with log rank test and Cox’s proportional hazards model was used to evaluate the overall survival (OS) since tumor resection between a high and low patient group of ImmuNEO patients. For continuous parameters, groups were divided by the median into high (above median) and low (below median) groups. For relative parameters (0-100%), patients were divided into a high group with fractions above 50% and low group with fractions below 50%. Here, only one representative tumor sample from each patient (ImmuNEO core cohort see Suppl. Table S1A) has been used to circumvent bias due to multiple metastasis available for some patients.

## Supplemental Information

**Supplementary Figure S1.**
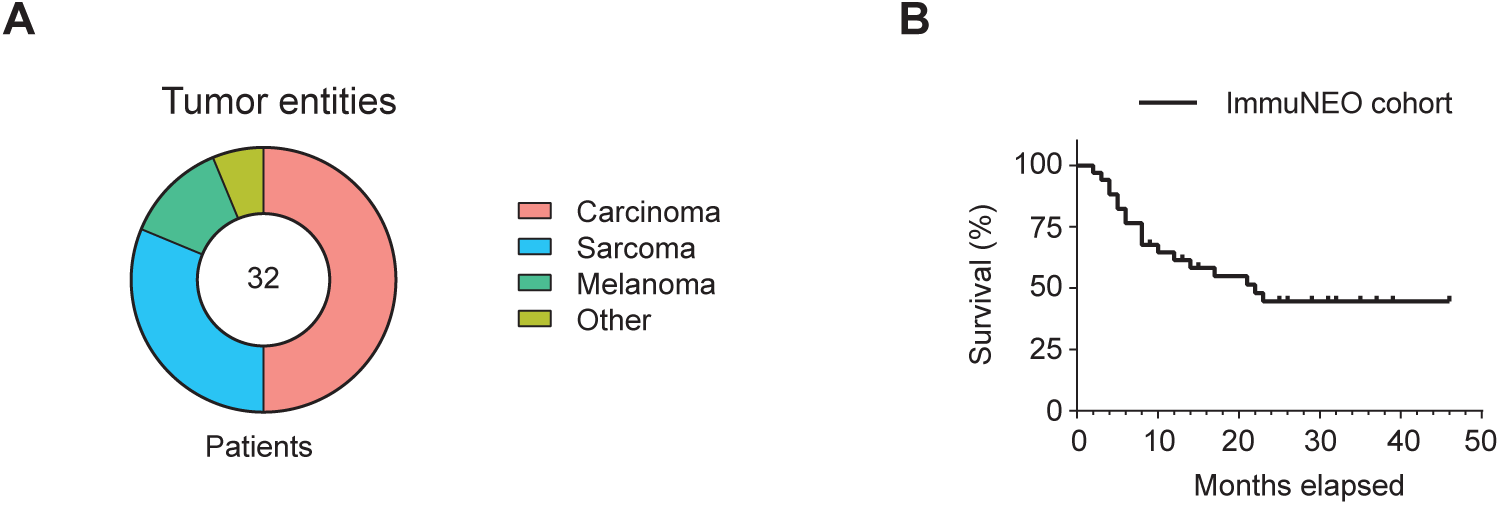
ImmuNEO MASTER cohort. **A,** Distribution of the major groups of tumor entities of patients included in the ImmuNEO MASTER cohort. **B,** Overall survival of ImmuNEO MASTER patients since tumor resection in months. **A, B,** n = 32 patients (see Suppl. Table S1A).

**Supplementary Figure S2.**
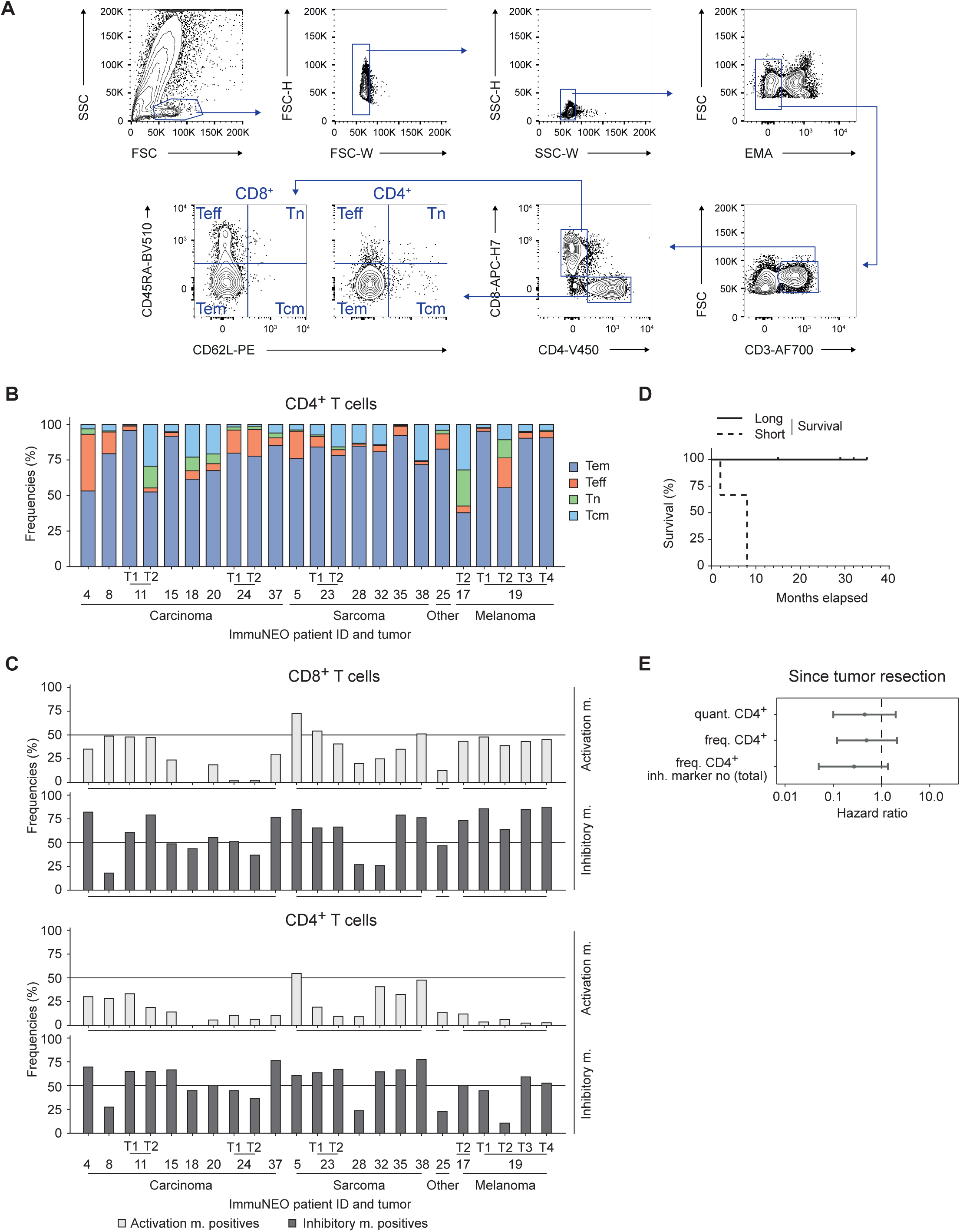
Analysis of the tumor microenvironment. **A,** Flow cytometric gating strategy for CD4^+^ and CD8^+^ T cell subsets. **B,** Frequencies of different CD4^+^ T cell subsets of all identified tumor-infiltrating CD4^+^ T cells per patient and grouped by tumor entity. **C,** Frequencies of CD4^+^ (bottom) and CD8^+^ T cells (top) per patient and grouped by tumor entity, expressing at least one activation marker (HLA-DR, CD103) or inhibitory marker (PD-1, TIM-3, LAG-3). **D,** Kaplan-Meier survival estimation since tumor resection of patients with short survival (below 1 year, n = 3) and long survival (above 1 year, n = 5). **E,** Forest plot showing the hazard ratio (dot) and 95% confidence intervals (lines) calculated by log rank test and Cox’s proportional hazards model of several phenotypic parameters for the survival of patients since tumor resection (n = 17). For statistical analysis only one representative tumor sample per patient was used (see core cohort Suppl. Table S1A). **A-C,** n = 23 tumor samples from n = 17 patients (see Suppl. Table S1A). freq., frequency; m., marker; NK, natural killer; quant., quantified cells per gram tumor; T, tumor; Tcm, central memory T cells; Teff, effector T cells; Tem, effector memory T cells; Tn, naïve T cells.

**Supplementary Figure S3.**
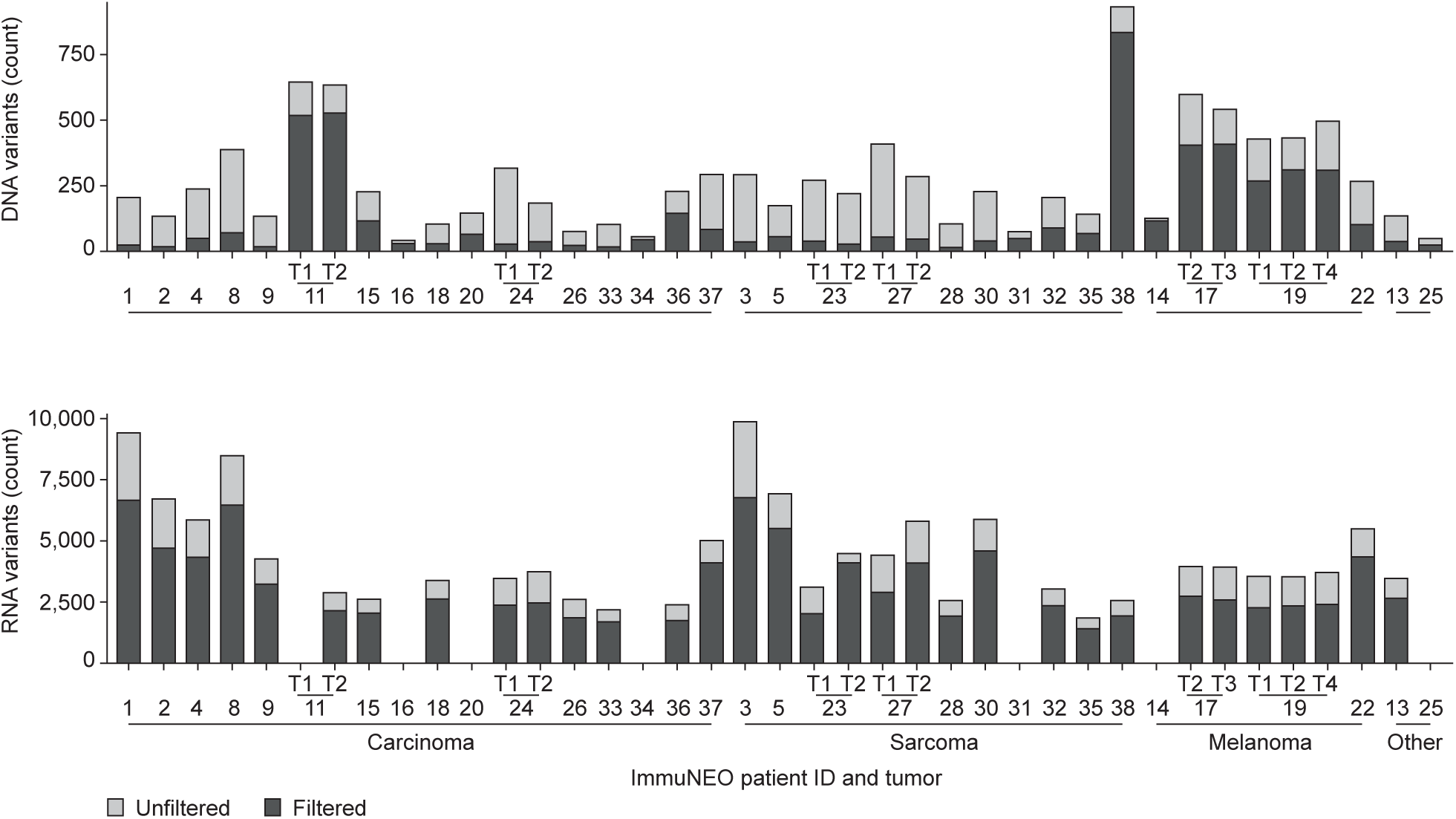
Quality assessment of genetic variants identified at the DNA and RNA level. Total unfiltered genetic variants identified by MuTect2 (v4.1.0.0) from whole exome (WES)/whole genome sequencing (WGS) data (DNA variants; upper panel) and by Strelka2 (v2.9.10) from RNA sequencing (RNA-seq) data (RNA variants; lower panel) are shown per tumor sample and grouped by tumor entity. Variants passed filtering for quality assessment only if they showed at least a coverage of 5 reads, a variant frequency of 5%, and 2 mutated reads within the tumor as well as not more than 1 mutated read within normal control tissue. n = 39 tumor samples from n = 32 patients for WES/WGS data; n = 32 tumor samples from n = 26 patients for RNA-seq data (see Suppl. Table S1A). T, tumor.

**Supplementary Figure S4.**
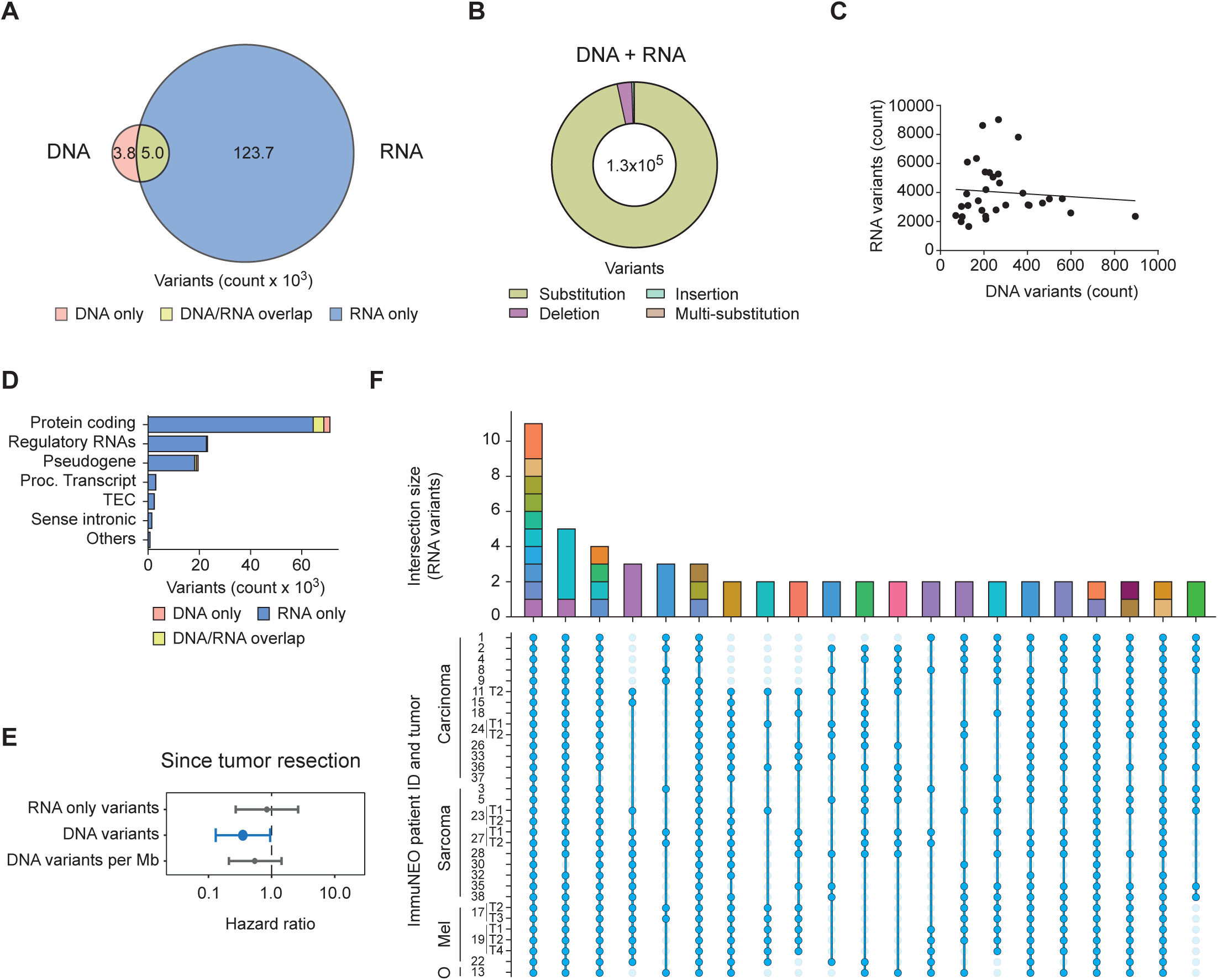
Genetic variants identified at the DNA and RNA level in tumor tissue from different cancer entities. **A,** Venn diagram showing the overlap between all variants identified from whole exome (WES)/whole genome sequencing (WGS) data (DNA variants) and from RNA sequencing (RNA-seq) data (RNA variants). **B,** Distribution of each mutation type for all identified genetic variants regardless of the sequencing origin (WES/WGS and RNA-seq combined). **C,** Correlation of DNA variants with RNA variants identified from tumor samples where matching WES/WGS and RNA-seq data was available (n = 32 tumor samples). Symbols depict individual tumor samples; Spearman’s rank correlation analysis, ρ = 0.1578; line depicts linear regression, R²=0.008. **D,** Bar graph showing the number of variants found in each genetic biotype and the originating dataset. **E,** Forest plot showing the hazard ratio (dot) and 95% confidence intervals (lines) calculated by log rank test and Cox’s proportional hazards model of several genetic parameters for the survival of patients since tumor resection (DNA variants n = 32 patients, RNA variants n = 26 patients). Significant results (p ≤ 0.05) are highlighted in blue. For statistical analysis only one representative tumor sample per patient was used (see core cohort Suppl. Table S1A). **F,** Upset plot showing the overlap of at least two RNA variants between at least ten tumor samples. The bar graph shows the number of unique variants present in a shared subset of tumors defined as intersection size, dots indicate the tumor samples where the subset is present, and lines connect tumor samples within the same subset. The different genes harbouring the defined genetic variants are coloured in the intersection bar graph. **A, B, D, F,** n = 32 tumor samples from n = 26 patients for WES/WGS data and for RNA-seq data (see Suppl. Table S1A). Mel, melanoma; Mb, mega base; O, other; OS, overall survival; Proc., processed; T, tumor; TEC, to be experimentally confirmed.

**Supplementary Figure S5.**
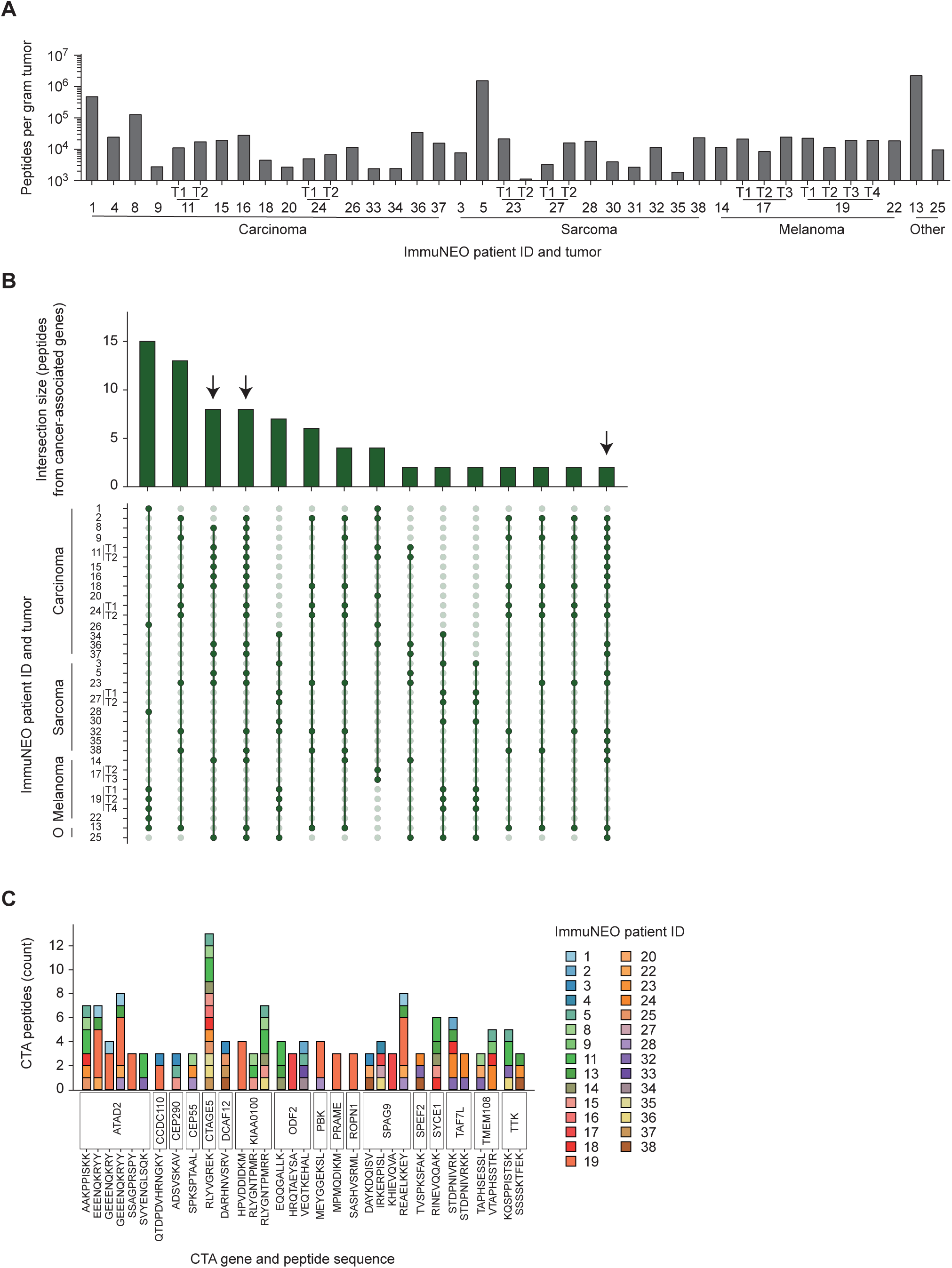
Characterization of shared cancer-associated peptides in the HLA class I immunopeptidomes. **A,** Quantitative numbers of unique HLA class I peptides per gram tumor identified per tumor sample grouped by tumor entity. Peptides bound to HLA class I molecules on the surface of tumor cells have been isolated by immunoprecipitation and sequenced by liquid chromatography with tandem mass spectrometry (LC-MS/MS). Peptide sequences were then mapped with 1% FDR to the Ensemble92 protein database using pFIND (v3.1.5) (23) and unique sequences have been filtered. **B,** Upset plot showing peptide overlap from tumor-associated genes defined by the Protein Atlas between all tumor samples. The bar graph shows the number of unique peptides present in a shared subset of tumors defined as intersection size, dots indicate the tumor samples where the subset is present, and lines connect tumor samples within the same subset. Subsets that present in at least 11 patients are highlighted by arrows. **C,** Bar graph showing peptides origination from the annotated cancer testis antigen (CTA) genes, their sequence, and the patients where they have been found. **A-C,** n = 41 tumor samples from n = 32 patients (see Suppl. Table S1A). CTA, cancer testis antigen; T, tumor.

**Supplementary Figure S6.**
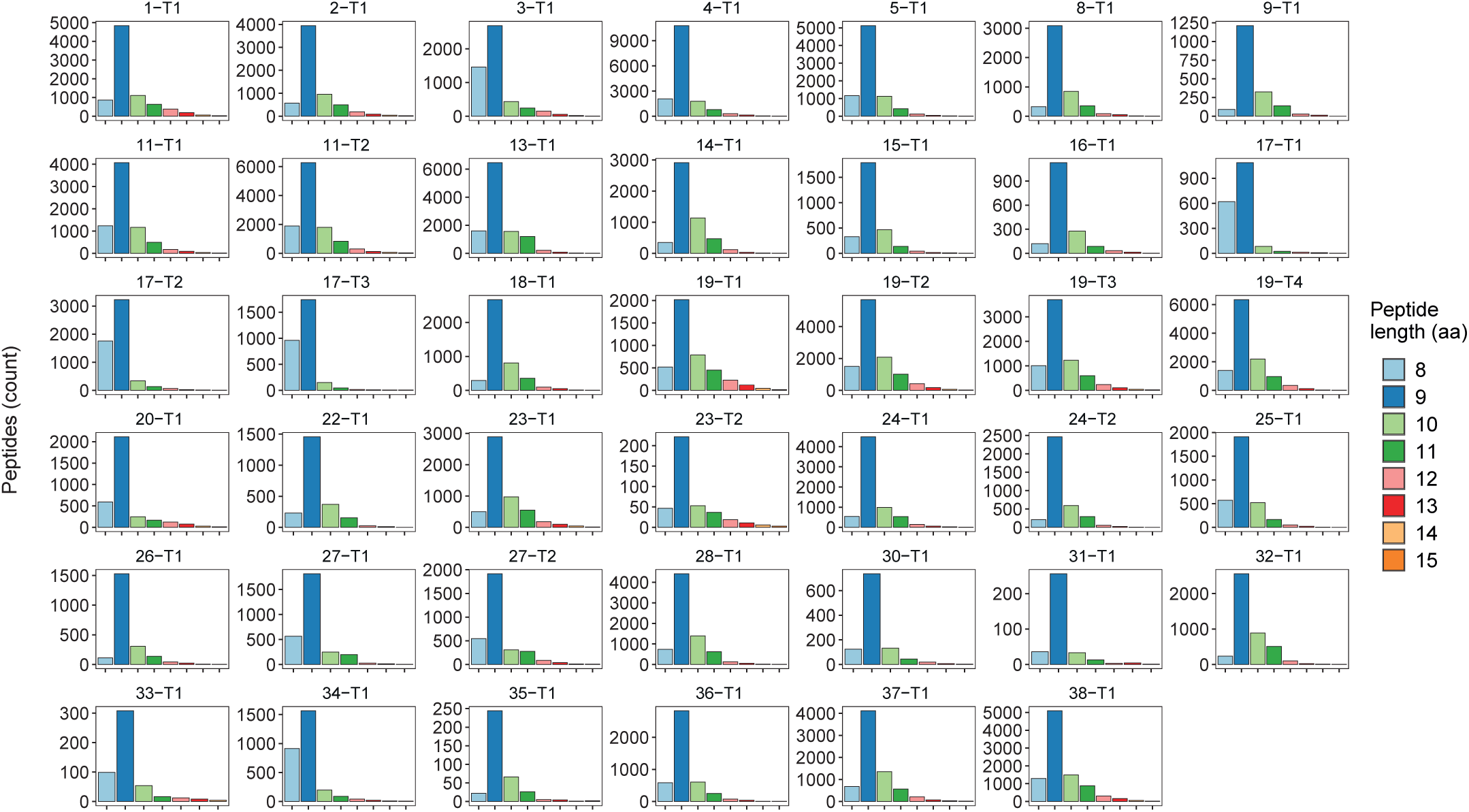
Length distribution of HLA class I peptides identified by mass spectrometry. Bar graph showing the number of unique peptides per peptide length in amino acids for every analysed tumor sample. Peptides bound to HLA class I molecules on the surface of the tumor cells have been isolated by immunoprecipitation and sequenced by liquid chromatography with tandem mass spectrometry (LC-MS/MS). Peptide sequences were then mapped with 1% FDR to the Ensemble92 protein database using pFIND (v3.1.5) and unique sequences have been filtered. n = 41 tumor samples from n = 32 patients (see Suppl. Table S1A). aa, amino acids; T, tumor.

**Supplementary Figure S7.**
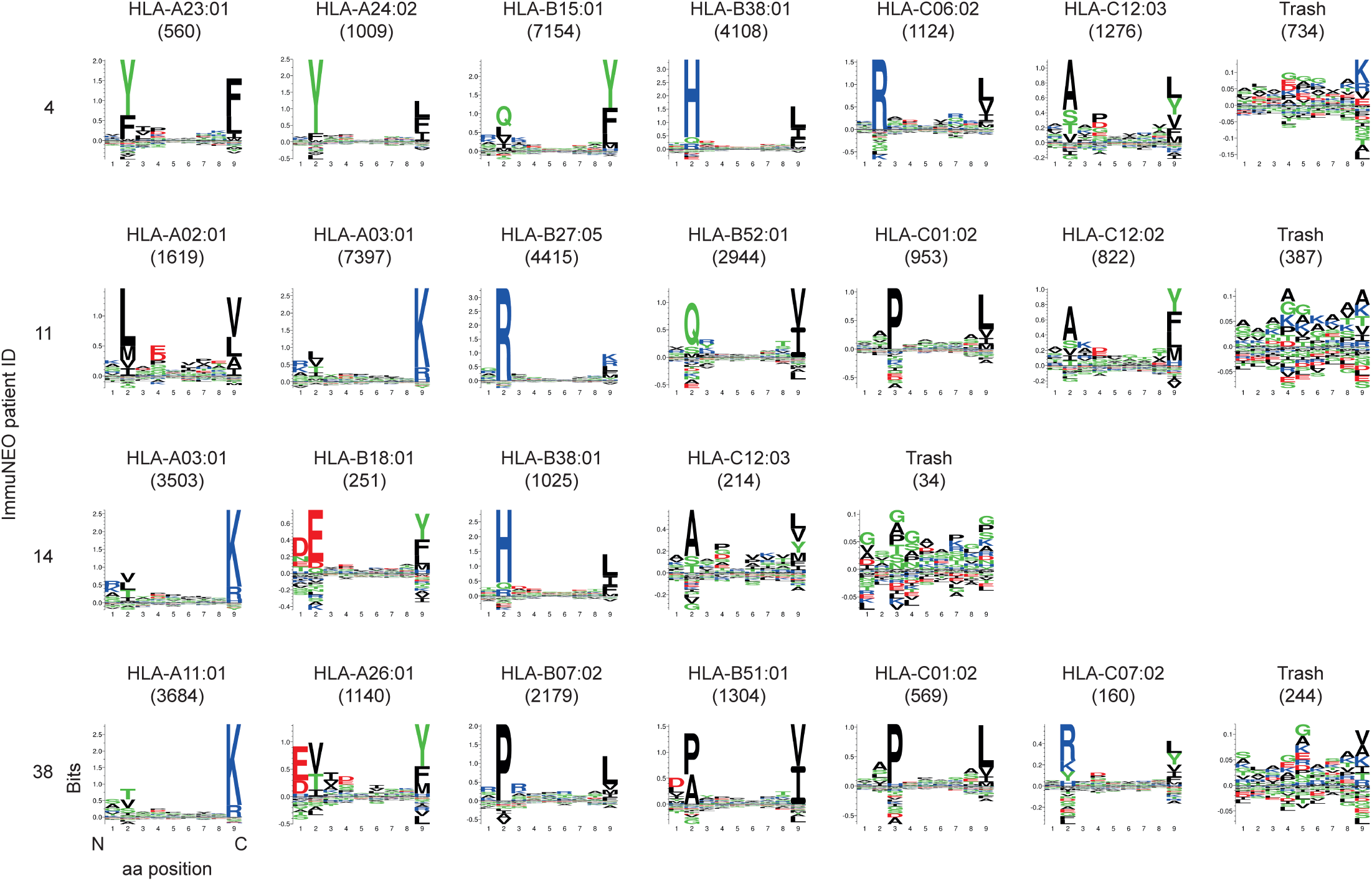
Peptide HLA class I binding motifs within the immunopeptidome. MHCMotifDecon (v1.0) has been used to match all isolated HLA class I peptides with lengths from 8- 15 amino acids to the patients’ HLA class I alleles according to their binding motifs and anchor residues for each tumor sample using standard settings. Binding motives of four representative tumor samples for each HLA class I allele are displayed with the total number of matched peptide sequences in brackets. Peptides not matching any HLA class I allele of the respective patient are displayed in the trash subgraph. aa, amino acid; HLA, human leukocyte antigen.

**Supplementary Figure S8.**
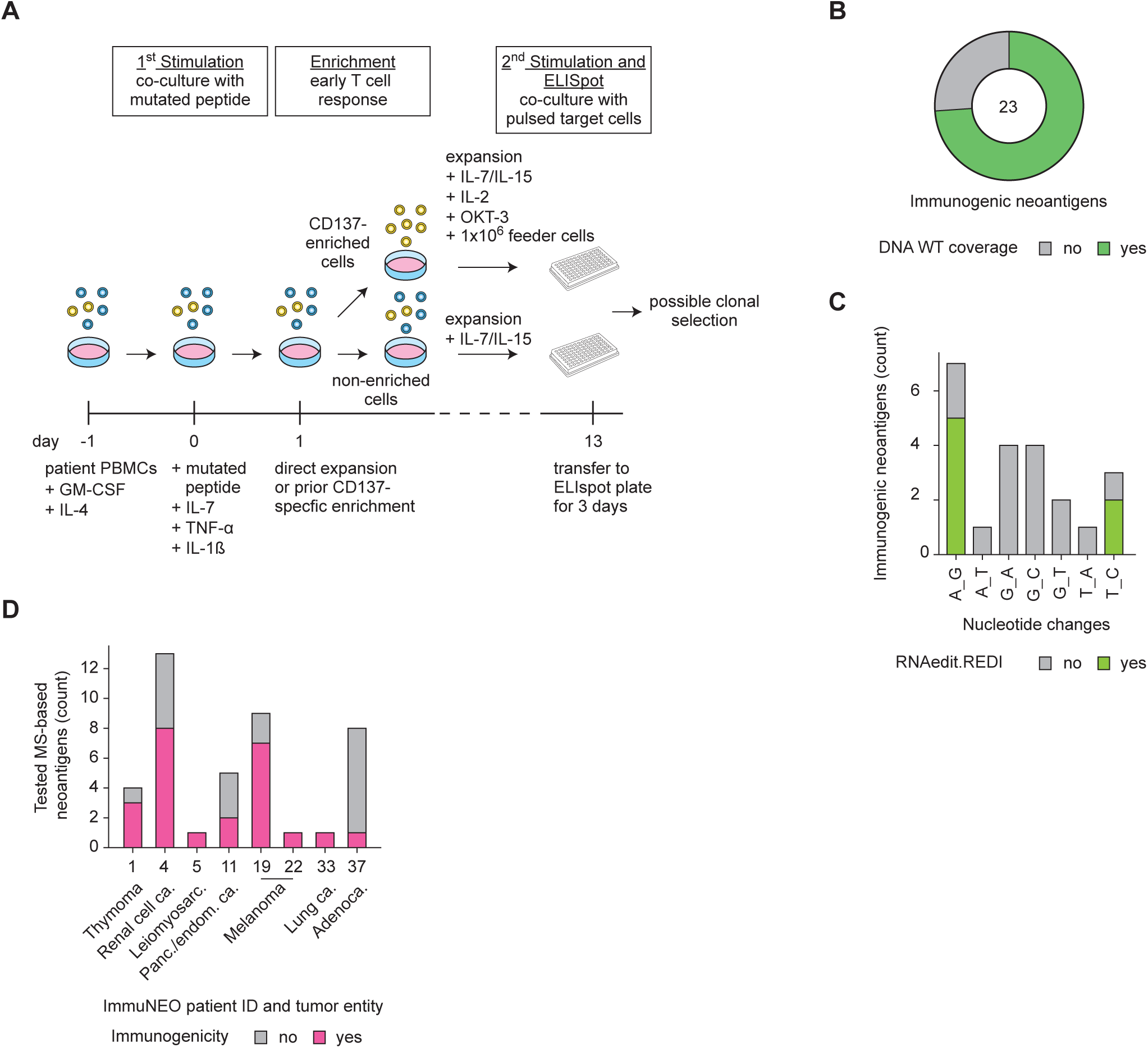
Characterization and immunogenicity assessment of neoantigen candidates. **A,** Schematic overview of the immunogenicity assessment by modified accelerated co-cultured dendritic cell (acDC) assay using non-enriched or CD137^+^-enriched T cells (PBMCs or TILs) for subsequent IFN-γ ELISpot readout. **B,** Pie chart depicting the proportion of immunogenic neoantigens identified only from RNA sequencing (RNA-seq) data where the respective wild type (WT) sequence was identified at the DNA level with a coverage of ≥ 3 reads (green) or the respective region was not covered at the DNA level (grey, < 3 reads). **C,** Distribution of the nucleotide exchange pattern of all single nucleotide variants that yield immunogenic neoantigen candidates identified only from RNA-seq data (n=22). Variants previously identified in the REDIportal (29) database as RNA editing events are highlighted in green. **D,** For those patients with immunogenic neoantigens, the total number of tested neoantigen candidates is depicted including immunogenic and non-immunogenic ones. **B-D,** n = 79 neoantigen candidates from n = 24 patients were analysed in total; n = 8 patients harboured n = 24 immunogenic neoantigens; n = 1 immunogenic neoantigens from DNA and RNA variants; n = 23 neoantigen candidates from RNA variants. ca., carcinoma; endom., endometrium; GM-CSF, granulocyte macrophage-colony stimulating factor; IL, interleukin; OKT-3, Muromonab-CD3; Panc., pancreas; TNF-α, tumor necrosis factor-α.

**Supplementary Table S1 Overview of the ImmuNEO patient cohort A,** Detailed information on every tumor sample of the ImmuNEO cohort including entity, metastatic site (or primary), stage at admission and primary sampling cohort. Core samples used for statistical analysis (subset) are labelled. Tumor samples used for the immune phenotyping of the tumor microenvironment (TME) by flow cytometric assessment and RNA sequencing (RNA-seq) of sorted CD8^+^ T cells are marked. Samples where whole exome sequencing (WES) and bulk tumor RNA-seq was performed are annotated; samples analysed via whole genome sequencing (WGS) are marked with an asterisk. The survival status as well as the survival times in months are displayed for several periods since initial diagnosis (ID), diagnosis of metastatic disease (MD) and since admission to MASTER/tumor resection (MASTER). The time difference since MD and MASTER is shown in months. Furthermore, information is given on patients receiving immune checkpoint blockade in general, prior to and after study admission and the respective response with no response (0), mixed response (1) and good response (2). **B,** Information about applied therapies for every ImmuNEO patient prior to and after tumor resection. 1 = therapy applied, 0 = therapy not applied. **C,** Table providing information on HLA class I alleles identified for each patient from whole exome (WES)/whole genome sequencing (WGS) data using the combination of the algorithms xHLA, BWAKit, and OptiType. For ImmuNEO-1, -4, -19 and -22 (*) the alleles were confirmed using targeted NGS (Zentrum für Humangenetik und Laboratoriumsdiagnostik, Martinsried, Germany). Ca, carcinoma; Chemo, chemotherapy; DSRCT, desmoplastic small round cell tumor; GIST, gastrointestinal stromal tumor; HLA, human leukocyte antigen; IME, immune microenvironment; IN, ImmuNEO; LN, lymph node; MPNST, malignant peripheral nerve sheath tumor; MS, mass spectrometry; OP, operation; RNAseq, RNA sequencing; WES, whole exome sequencing; WGS, whole genome sequencing.

**Supplementary Table S2 Genetic variants A, B, C,** Table showing the somatic variants present in at least four patients **(A)**, the RNA alterations present in min. 10 patients **(B)** and the RNA alterations (min. 2 in each group) present in min. 10 unique samples **(C)**. Variants were called by MuTect2 (v4.1.0.0) from whole exome (WES)/whole genome sequencing (WGS) data and by Strelka2 (v2.9.10) from RNA sequencing (RNA-seq) data. SNP-filtering has been performed using the dbSNP-all data base. No RNA data was available for patients IN-11-T1, IN-14, IN-16, IN-20, IN-25, IN-31, IN-34. For every variant (Mutation_ID consist of chromosome, position, reference base and alternative base) the affected gene and the gene biotype are shown. The samples and patients where the variant is present is shown and counted (upper table). In the lower table all information for each variant within each sample is listed. The number of wt reads (TumorRD) and mutated reads (TumorAD) in the tumor, the wt reads (NormalRD) and mutated reads (NormalAD) in the normal control, the variant frequency within the tumor (TumorVF) and the coverage of each mutation within the tumor (TumorCoverage) is included. IN, ImmuNEO.

**Supplementary Table S3 Tumor-associated peptides shared between several tumor samples** Table shows the sequence of wild-type peptides found in at least eleven tumor samples from tumor-associated genes defined by the ProteinAtlas. Peptides bound to HLA class I molecules on the surface of tumor cells have been isolated by immunoprecipitation and sequenced by liquid chromatography with tandem mass spectrometry (LC-MS/MS). Peptide sequences were then mapped with 1% FDR to the Ensemble92 protein database using pFIND (v3.1.5) and unique sequences have been filtered. The MS count and the length of each peptide in amino acids (aa) is annotated as well as the predicted binding affinity and rank for two HLA-I alleles present in all patients defined by NetMHC4.0. aa, amino acids; FDR, false discovery rate; HLA, human leukocyte antigen; IN, ImmuNEO; nM, nanomol; SB, strong binder; Seq, sequence; WB, weak binder.

**Supplementary Table S4 Overview of mutated peptide ligands A,** Detailed information on all neoantigen candidates. By combining genomic mutational data with mass-spectrometry (MS)-based immunopeptidomic data for each patient sample, neoantigen candidates have been identified. pFIND (v3.1.5) was used at 5% FDR on spectral level for the identification of non-wild type (WT) 8-15mer neoantigen candidates. The machine learning tool Prosit was additionally integrated to rescore and rematch the peptide spectra using unfiltered pFIND data as input. n = 39 tumor samples from n = 32 patients were analysed in total; n = 27 tumor samples from n = 24 patients harboured n = 91 neoantigen candidates. Using netMHC4.0 and MHCFlurry, binding predictions for each peptide towards the patients six HLA class I alleles has been performed and for each algorithm the best binding allele by affinity and by rank are shown. Mutated amino acids are marked with two asterisks within the sequence. **B,** Additional information on immunogenic neoantigen candidates. Immunogenicity assessment has been performed using a modified accelerated co-cultured dendritic cell (acDC) assays with IFN-γ ELIspot analysis using patient derived PBMC (non-enriched and CD137^+^ enriched) or tumor-infiltration lymphocytes (TILs) (non-enriched and CD137^+^ enriched) (**top table**) and allogenic-matched healthy donor PBMCs (non-enriched) (**bottom table**). Shown are immunogenic neoantigens that elicit an immune response where the ratio of spot forming units (SFU) is > 2 (mutated / control peptide) and the difference of SFU is > 50 (mutated - control peptide). Autologous lymphoblastoid cell lines (LCLs) or allogenic HLA-matched cells (LCLs or HLA-transduced cell lines) have been used as target cells. a.a, amino acid; Alt, alternative; BA, binding affinity; CA, carcinoma; Chrom, chromosome; del, deletion; dup, duplication; HD, healthy donor; HLA, human leukocyte antigen; ins, insertion; n.a./NA, not applicable; nM nanomole; PBMC, peripheral blood mononbuclear cells; Pos, position; Ref, reference; Seq, sequence; T, tumor; VF, variant frequency.

## Acknowledgements

We thank G. Swinerd and S. Mall for the preliminary organization of sample collection, the measurement of some patient samples and the preliminary assembly of flow-cytometry antibody panels. We thank the virology department of MRI of U. Protzer for technical support and advice with Elispot measurements and analysis. We especially thank A. Stelzl for logistics organization, measurement of patient samples and experimental support. We thank the NCT Molecular Precision Oncology Program for technical support and funding through project number 21.

